# Pancreatic tumor microenvironment reprogramming via alloantigen-expressing virotherapy elicits tumor rejection and improves immunotherapy response

**DOI:** 10.1101/2025.09.15.675641

**Authors:** Mulu Z. Tesfay, Aleksandra Cios, Zetao Cheng, Khandoker U. Ferdous, Randal S. Shelton, Bahaa Mustafa, Natalie M. Elliott, Elizabeth A. Raupach, Camila C. Simoes, Isabelle R. Miousse, Alicja Urbaniak, Michael A. Bauer, Eric R. Siegel, Steven R. Post, Jean C. Chamcheu, Rang Govindarajan, Valery Z. Grdzelishvili, Martin J. Cannon, Martin E Fernandez-Zapico, Alexei G. Basnakian, Chiswili Y. Chabu, Omeed Moaven, Mitesh J. Borad, Bolni M. Nagalo

**Author notes:** **Correspondence**: Bolni Marius Nagalo, BSc., MSc., Ph.D. Associate Professor Program in Oncology, Department of Pharmacology and Physiology, University of Maryland School of Medicine, Marlene and Stewart Greenebaum NCI Comprehensive Cancer Center, University of Maryland School of Medicine, 655 W. Baltimore St., Baltimore, MD, 21201, USA,; Mitesh J. Borad, MD Professor of Medicine, Division of Hematology and Medical Oncology, Mayo Clinic in Arizona, Mayo Clinic Comprehensive Cancer Center, Mayo Clinic, 5777 East Mayo Boulevard, Phoenix, AZ 85254, USA. These authors contributed equally to this work (Mulu Z. Tesfay, Aleksandra Cios).

## Abstract

Pancreatic ductal adenocarcinoma (PDAC), the most common malignant type of pancreatic cancer, is characterized by a dense desmoplastic stroma, low neoantigen burden, and a highly immunosuppressive tumor microenvironment (TME), which severely limit cytotoxic T-cell infiltration and the efficacy of immune therapies. Here, we present a novel strategy harnessing acute transplant rejection mechanisms by employing a recombinant oncolytic rVMG vector engineered to express the murine H-2Kk MHC class I alloantigen (rVMG-H-2Kk), thereby inducing tumor-specific antigenic mismatch responses. *In vitro*, rVMG-H-2Kk exhibited strong replication and cytolytic activity while inducing cell surface expression of both H-2Kk and endogenous H-2Kb, in addition to upregulation of antigen presentation genes (β2-microglobulin, Tap1, and Tapbp). In two independent immunocompetent PDAC models, intratumoral and systemic delivery of rVMG-H-2Kk delayed tumor progression and prolonged survival. Multiplex immunohistochemistry and immunophenotyping revealed substantial TME remodeling, marked by increased effector T-cell infiltration, regulatory T-cell depletion, and reduced fibrosis. Spatial transcriptomics further showed compartment-specific immune activation and epithelial metabolic reprogramming, corroborating with enhanced tumor immunogenicity. Despite these effects, rVMG-H-2Kk also induced compensatory immunosuppressive pathways, including upregulation of antiviral response genes and immune checkpoint receptors such as PDL-2. Importantly, combination therapy with rVMG-H-2Kk and murine checkpoint blockade (anti-PD-1 and anti-CTLA-4) drastically improved survival than checkpoint blockade alone. Strikingly, surviving mice resisted tumor rechallenge, indicating the establishment of durable antitumor memory. Collectively, these findings establish rVMG-H-2Kk as a novel immunotherapeutic platform capable of converting immune-cold tumors into immune-hot, sensitizing tumors to immune checkpoint inhibitors, and establishing durable antitumor immunity in PDAC.

**One Sentence Summary:** Oncolytic virus-based delivery of an alloantigen improves antitumor immunity and synergizes with immune checkpoint inhibitors in pancreatic cancer.

**Significance statement:** Delivery of murine alloantigens via an engineered oncolytic vesiculovirus, combined with immune checkpoint blockade, overcomes immunosuppressive barriers and establishes durable antitumor immunity in PDAC.

## INTRODUCTION

Pancreatic ductal adenocarcinoma (PDAC), which accounts for approximately 85% of all malignant pancreatic cancer cases, remains among the deadliest malignancies, with limited therapeutic options and poor long-term survival.(1) Its immunologic landscape is characterized by a dense desmoplastic stroma and a profoundly immunosuppressive tumor microenvironment (TME), which restrict cytotoxic T lymphocyte (CTL) infiltration and impairs the efficacy of immune therapies. (2–5) The TME in PDAC is dominated by tumor-associated macrophages, myeloid-derived suppressor cells (MDSCs), and regulatory T cells, with scarce infiltration of functional tumor-reactive lymphocytes.(6) Compounding this immune exclusion, PDAC tumors often lack sufficient neoantigen expression to drive effective endogenous T-cell priming.(7) As a result, immunotherapeutic agents such as immune checkpoint inhibitors have shown limited benefit, with responses largely confined to microsatellite instability-high, mismatch repair-deficient, POLE-mutated, or homologous recombination-deficient tumors.(8–13) Since these molecular alterations are present in only a minority of cases, the overall responsiveness of PDAC to immunotherapy remains severely limited.

To address this challenge, we explored the immunologic principles of transplant rejection to enhance tumor-specific immune responses. Alloantigen mismatch, the immunologic foundation of transplant rejection, is one of the most potent triggers of host immune activation.(14) Rejection occurs in distinct stages—hyperacute (antibody- and complement-mediated), acute (T-cell–driven), and chronic (associated with fibrosis and vascular remodeling). (15) Although recent studies have leveraged xenogeneic antigens to stimulate alloimmune responses, (16, 17) the use of defined, immunologically relevant alloantigens remains largely underexplored. A first-in-human trial demonstrated that systemic delivery of Newcastle disease virus expressing the porcine α1,3-galactosyltransferase antigen-induced hyperacute rejection-like responses, achieving a 90% disease control rate without serious adverse events in patients with refractory metastatic cancers.(17)

Building on this rationale, we developed a chimeric oncolytic vesiculovirus platform (rVMG), in which the glycoprotein of VSV is replaced with that of Morreton virus to enhance tumor targeting and reduce neurotoxicity.(18) We further engineered this vector to express the murine MHC class I alloantigen H-2Kk (rVMG-H-2Kk), with the goal of triggering an alloantigen mismatch and redirecting host immune responses against the tumor. In vitro, rVMG-H-2Kk demonstrated robust replication and cytolytic activity in murine PDAC cells, along with strong surface expression of H-2Kk, upregulation of endogenous H-2Kb, and increased expression of major antigen presentation genes (β2-microglobulin, Tap1, and Tapbp).

Next, we evaluated rVMG-H-2Kk in two immunocompetent murine PDAC models. In both models, rVMG-H-2Kk reprogrammed the TME toward a proinflammatory state, delayed tumor growth, and extended survival compared to parental rVMG and PBS controls. In the KPC model, histopathologic analysis revealed reduced fibrosis and α-smooth muscle actin expression, suggesting stromal remodeling. Serum biochemistry confirmed an absence of severe toxicity, and spatial transcriptomics indicated immune activation alongside compensatory upregulation of immunosuppressive mediators such as PDL-2 and interferon-stimulated genes. To overcome this adaptive resistance, we combined systemic rVMG-H-2Kk with dual checkpoint blockade (anti– PD-1 and anti–CTLA-4), which significantly improved survival, with 27% of mice surviving to day 90. Importantly, rechallenged survivors rejected tumor regrowth and exhibited cytokine signatures consistent with durable antitumor memory. Together, these findings highlight the potential of rVMG-H-2Kk as a next-generation immunovirotherapy platform capable of reprogramming the PDAC immune landscape, sensitizing tumors to immune checkpoint inhibitors, and eliciting long-lasting antitumor immunity.

## RESULTS

### Engineered viral vector rVMG-H-2Kk retains replicative fitness and cytolytic activity in murine PDAC cells

We generated rVMG by replacing the VSV glycoprotein gene with that of Morreton virus. (18, 19) The rVMG-H-2Kk variant was engineered to express the murine H-2Kk alloantigen and generated using reverse genetics in BHK-21 cells as previously described (Fig. 1A).(18) In KPC and Pan02 cells infected at a multiplicity of infection (MOI) of 0.1, real-time impedance analysis showed a reduction in cell viability compared to mock-infected controls. Both rVMG and rVMG-H-2Kk induced similar cytopathic effects in tumor cells 72 hours post-infection (Fig. 1, B to C. An MTS assay demonstrated a significant, dose-dependent decrease in cell viability across all tested MOIs (10, 1, 0.1, and 0.01; fig. S1, A and B; *P* < 0.05). One-step growth assays in Vero cells showed that rVMG-H-2Kk replicated at levels comparable to the parental rVMG, with less than a 0.4-log difference in viral titers (Fig. 1, E and F), indicating that insertion of the alloantigen did not compromise viral fitness. Flow cytometric analysis demonstrated robust surface expression of endogenous H-2Kb in KPC cells infected with either rVMG (83.9%) or rVMG-H-2Kk (70.5%), suggesting a virus-mediated upregulation of MHC class I molecules (Fig. 1G, left panel). In parallel, rVMG-H-2Kk–infected cells showed high-level surface expression of the ectopic alloantigen H-2Kk (93.7%), while rVMG-infected cells exhibited only minimal H-2Kk signal (2.45%) (Fig. 1G, right panel), confirming efficient delivery and expression of the non-self MHC class I molecule. Moreover, qPCR analyses at 6 hours post-infection showed that VSV-N transcripts, which encode the nucleocapsid protein essential for viral replication, were upregulated by approximately 1,000-fold compared to mock-infected controls (*P* < 0.0001) (Fig. 1H), confirming active infection in rVMG-H-2Kk-infected cells. In addition, H-2Kk and endogenous MHC class I molecules (H-2Kb) transcripts were upregulated by ∼500x and ∼40x, respectively, in rVMG-H-2Kk-infected cells compared to mock-infected controls (Fig. 1, I and J). Furthermore, β2-microglobulin expression increased by approximately 50% in both treatment groups (rVMG, *P* = 0.0011; rVMG-H-2Kk, *P =* 0.0009; Fig. 1K), and Tap1 expression was upregulated by nearly 80% (rVMG, *P =* 0.0005; rVMG-H-2Kk, *P =* 0.0003; Fig. 1L). In contrast, Tapbp expression was significantly reduced in both groups—by ∼55% with rVMG and ∼43% with rVMG-H-2Kk (*P <* 0.0001 for both; Fig. 1M).

**Fig. 1.**
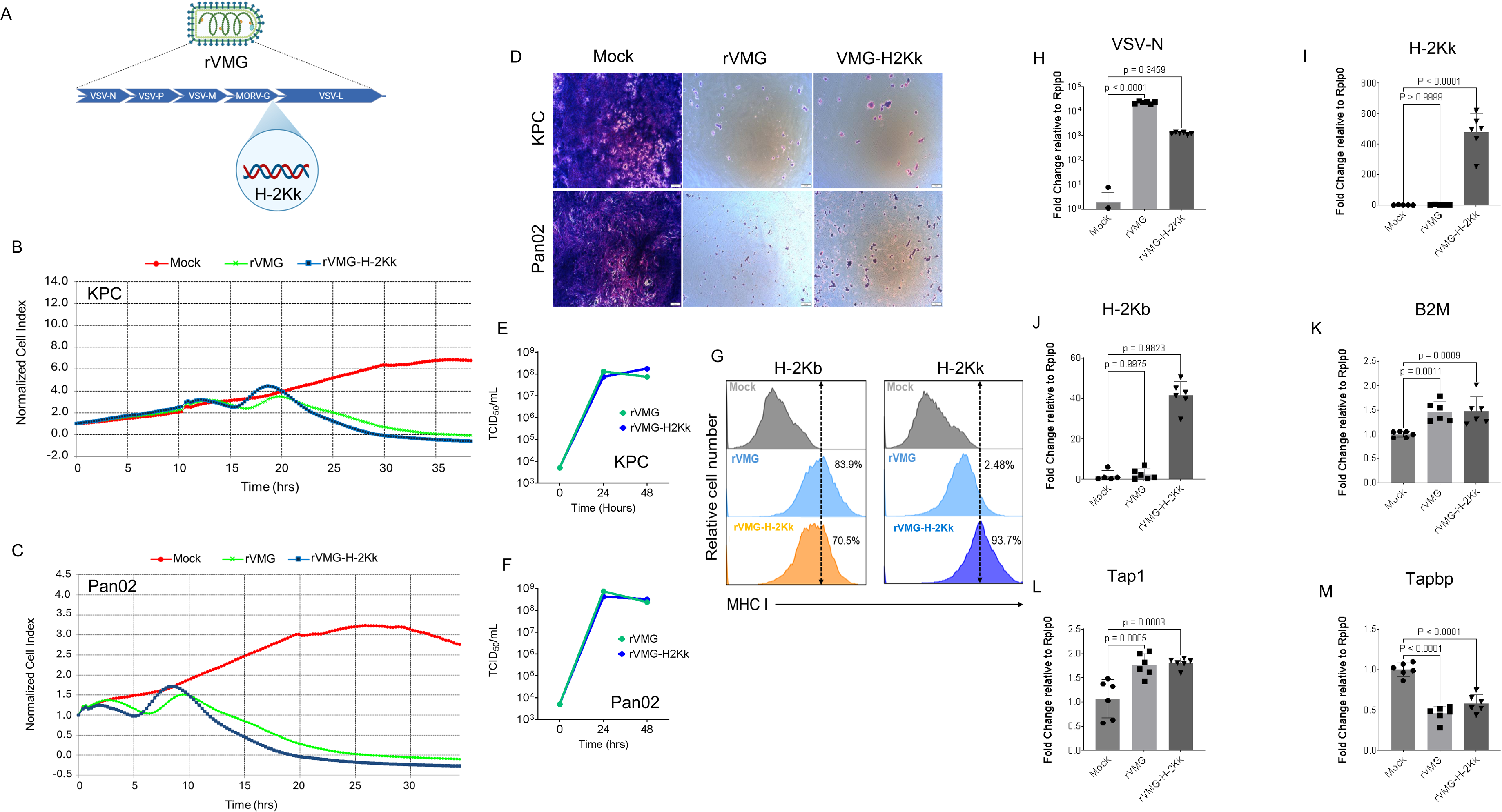
Characterization of rVMG-H-2Kk in murine pancreatic cancer cell lines. (**A**) Schematic of rVMG (top) and rVMG-H-2Kk (bottom) indicating the insertion of the H-2Kk alloantigen (P04223-HA1K_MOUSE H-2Kk). (**B** to **C**) Real-time impedance-based viability curves of KPC (**B**) and Pan02 (**C**) cells infected at a multiplicity of infection (MOI) of 0.1 with rVMG or rVMG-H-2Kk, or mock-infected, as measured by the xCELLigence system. (**D**) Brightfield images of KPC cells at 72 hours post-infection (hpi) in rVMG-H-2Kk–infected cells compared to mock-infected controls. (**E** to **F**) One-step viral growth assays in Vero cells infected at an MOI of 0.1, with viral titers determined in BHK-21 cells at 24 and 48 hpi. (**G**) Flow cytometry analysis of cell surface H-2Kb (left panel) and H-2Kk (right panel) expression in KPC cells following infection with rVMG or rVMG-H-2Kk. (**H** to **M**) Quantitative RT-PCR of VSV-N, H-2Kk, H-2Kb, B2M, Tap1, and Tapbp mRNA levels in KPC cells at 6 hpi (MOI = 0.1). Bars represent mean ± SEM from six samples. Statistical significance was determined by one-way ANOVA with *P* < 0.05 considered significant.

### Intratumoral rVMG-H-2Kk therapy delays tumor growth and improves survival in a subcutaneous Pan02 syngeneic model

Based on these indications that rVMG-H-2Kk infection upregulates genes involved in antigen processing, presentation, and immunoreactivity in cancer cells, we next sought to assess its effects in tumor-bearing animals. Immunocompetent mice with established subcutaneous Pan02 tumors received three weekly intratumoral (IT) injections of the vehicle (PBS), rVMG, or rVMG-H-2Kk (1 x 10^7^ TCID_50_) (Fig. 2A). Both rVMG (P = 0.0469) and rVMG-H-2Kk (*P* = 0.0004) significantly delayed tumor growth compared to PBS, with rVMG-H-2Kk achieving an estimated ∼69% reduction in endpoint tumor volume (Fig. 2, B to D). Kaplan Meier analysis in an independent cohort demonstrated that only rVMG-H-2Kk provided a significant survival benefit compared to PBS (*P* = 0.0027), while rVMG alone did not improve survival (*P* = 0.9999) (Fig. 2E). Body weight, which serves as a surrogate for toxicity, remained stable during treatment (fig. S1C).

**Fig. 2.**
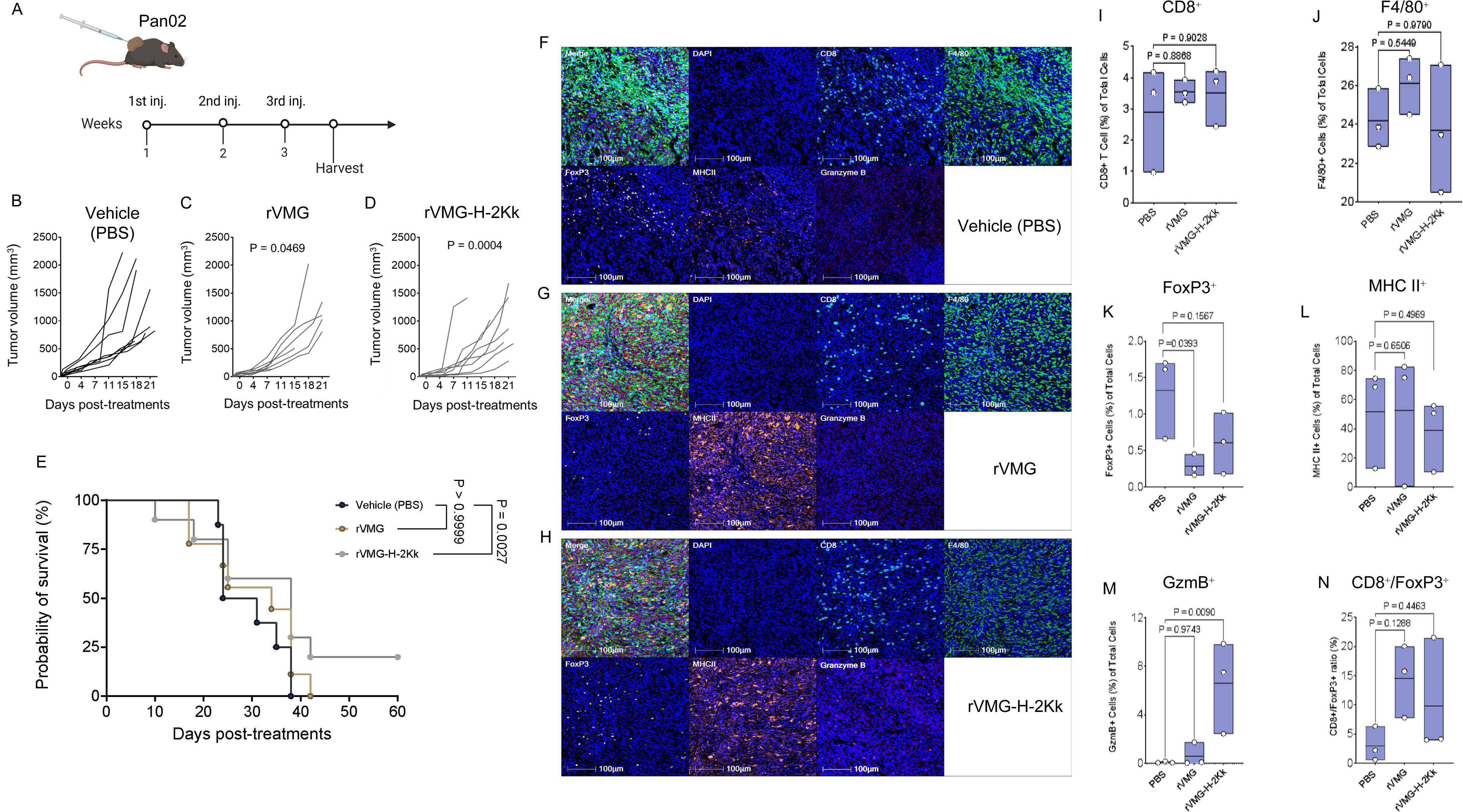
Therapeutic activity of intratumoral administration of rVMG-H-2Kk in subcutaneous Pan02 tumors. (**A**) Schematic of the treatment regimen showing three intratumoral injections of vehicle (PBS), rVMG, or rVMG-H-2Kk (1 x 10^7^ TCID_50_), with tumors harvested 3 days after final virus injection for immunoprofiling. (**B** to **D**) Individual tumor volume curves for each treatment group. (**E**) Kaplan–Meier survival analysis across treatment groups from an independent cohort. (**F** to **H**) Representative multiplex immunohistochemistry images of tumor-infiltrating lymphocytes in vehicle (PBS), rVMG, and rVMG-H-2Kk–treated tumors. (**I** to **N**) Quantification of CD8^+^, F4/80^+^, FoxP3^+^, MHC II^+^, and granzyme B^+^ (GzmB^+^) cells, and the CD8^+^/FoxP3^+^ ratio in the tumor microenvironment. Bars represent mean ± SEM from three tumors per group. Statistical significance was determined by one-way ANOVA with *P* < 0.05 considered significant.

### rVMG-H-2Kk reprograms the tumor microenvironment toward a proinflammatory state in Pan02 syngeneic tumors

Next, we employed multiplex immunohistochemistry (IHC) of the TME to assess immune cell subsets (list of antibodies in table S1). The levels of total CD8^+^ T cells, F4/80^+^ macrophages, and MHC II^+^ expression were similar among groups (Fig. 2, I, J, and L). However, rVMG treatment reduced the frequency of FoxP3 regulatory T cells by roughly 5-fold (∼5x) compared to PBS (*P* = 0.0393; Fig. 2K), while rVMG-H-2Kk treatment led to a marked increase in granzyme B (GzmB) T cells (*P* = 0.0090), increasing from ∼0% in PBS-treated tumors to ∼7% (Fig. 2M).

Flow cytometry analysis was performed to further characterize tumor-infiltrating lymphocytes (list of antibodies in table S2). We analyzed the distribution of key T cell subsets within the tumor microenvironment, including total CD4 and CD8 T cells, CD8 GzmB cytotoxic T cells, CD8 PD-1 exhausted T cells, and T follicular helper (Tfh) cells (fig. S2A–H). Notably, the frequency of CD8 T cells was nearly doubled in the rVMG group compared to PBS (*P* = 0.0023; fig. S2E), while rVMG-H-2Kk–treated tumors showed a substantial (>50x) reduction in exhausted CD8 PD-1 T cells compared to the PBS and rVMG groups (*P* = 0.0095; fig. S2G). We also evaluated the myeloid compartment and innate immune populations, including macrophages, dendritic cells (DCs), myeloid-derived suppressor cells (MDSCs), and natural killer (NK) cells (Fig. 3A–D). Myeloid cell profiling revealed that rVMG-H-2Kk reduced the total macrophage population by >3x relative to PBS (*P* = 0.0173; Fig. 3E). Both rVMG and rVMG-H-2Kk increased infiltration of pro-inflammatory M1 macrophages by approximately 1.5-fold (Fig. 3F) and decreased the frequency of anti-inflammatory M2 macrophages by >5x (*P* = 0.0274 for rVMG; *P* = 0.0367 for rVMG-H-2Kk; Fig. 3G). As a result, the M1/M2 ratio was elevated in both treatment groups (Fig. 3H). Granulocytic myeloid-derived suppressor cells (gMDSCs) were increased following both treatments; however, the proportion of gMDSCs in rVMG-treated tumors was highly variable (mean ± SD: ∼17% ± 20%), whereas rVMG-H-2Kk treatment resulted in a more consistent and pronounced increase (mean ± SD: ∼30% ± 6%). The ∼7.5-fold increase in gMDSCs relative to PBS-treated controls (mean ± SD: ∼4% ± 2%) was statistically significant (P = 0.0174; Fig. 3K). In contrast, monocytic MDSCs (mMDSCs) were significantly reduced by ∼4.0-fold in rVMG-treated tumors (P = 0.0228) and by ∼3.5-fold following rVMG-H-2K^k treatment (P = 0.0002; Fig. 3L). Finally, DCs and natural killer (NK) cells remained unchanged (Fig. 3, I to J).

**Fig. 3.**
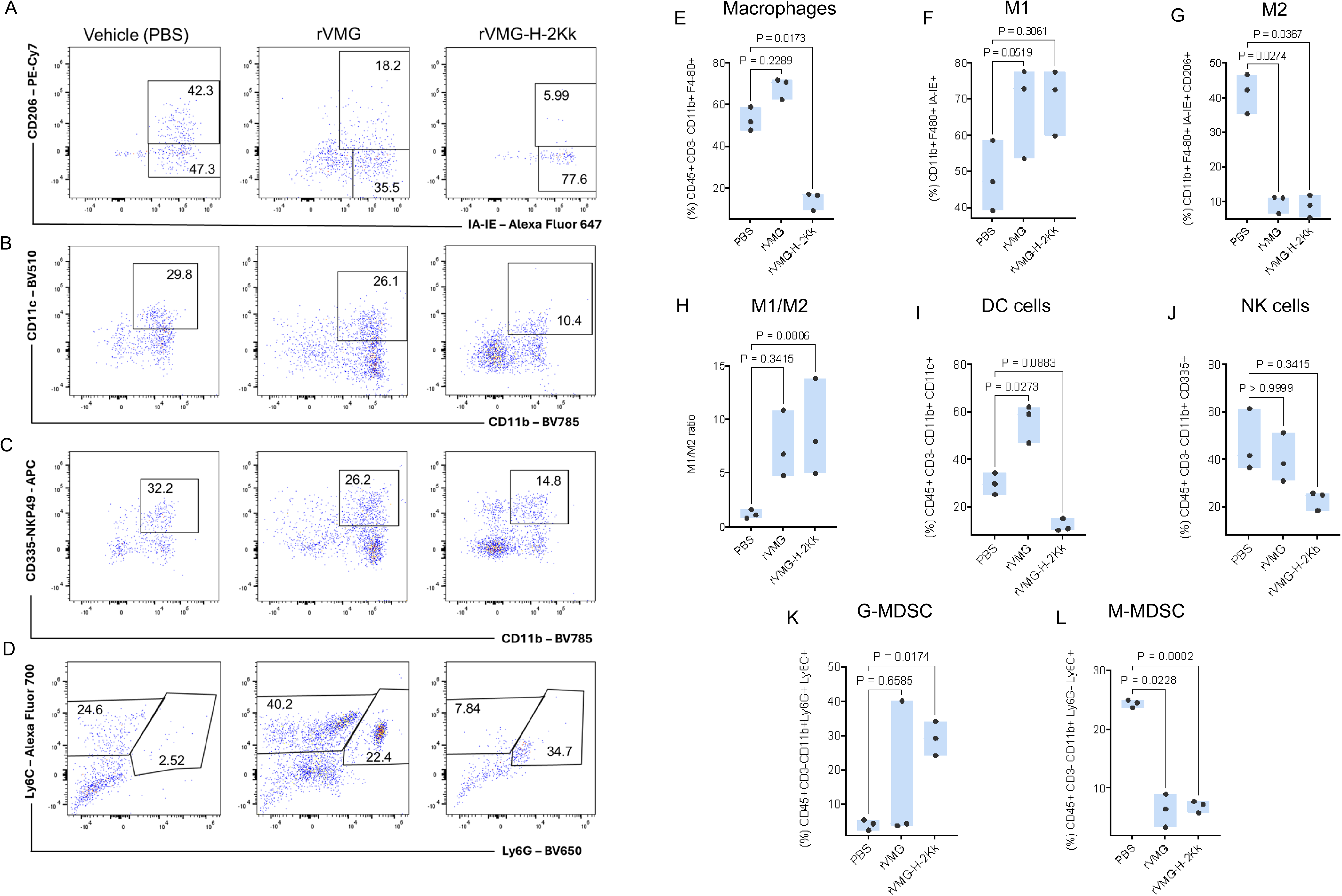
Flow cytometry gating strategies and immune subset distribution in subcutaneous Pan02 tumors. (**A**) Representative gating strategy for CD8^+^ T cells. (**B**) Gating strategy for regulatory T cells (Tregs; CD4^+^FoxP3^+^). (**C**) Gating strategy for M1- and M2-like macrophages within the tumor microenvironment. (**D** to **I**) Quantification of total T cells, CD4^+^ T cells, CD8^+^ T cells, CD8^+^GZmb^+^ T cells, CD8^+^ PD-1^+^ T cells, and Tfh cells across different treatment groups. Bars represent mean ± SEM from three tumors per group. Statistical significance was determined by one-way ANOVA with *P* < 0.05 considered significant. DC, dendritic cell; NK, natural killer; MDSC, myeloid-derived suppressor cells; G-MDSC, granulocytic MDSC; M-MDSC, monocytic MDSC

### Systemic delivery of rVMG-H-2Kk improves survival and remodels the TME in an orthotopic KPC model

To model metastatic disease amenable to systemic therapy, we evaluated the efficacy of rVMG-H-2Kk in an orthotopic KPC model, focusing on its impact on survival and its capacity to traffic to the tumor site to induce proinflammatory remodeling of the TME. Specifically, we assessed the immunomodulatory potential of rVMG and rVMG-H-2Kk vectors by examining intratumoral bioavailability, immune modulation, and survival outcomes. Mice bearing surgically implanted KPC tumors received three weekly intraperitoneal injections (100 µL each) of the vehicle (PBS), rVMG, or rVMG-H-2Kk (1 x 10^7^ TCID_50_) (Fig. 4A). Treatment with rVMG-H-2Kk, but not rVMG, conferred a statistically significant survival advantage over PBS (Vehicle), as determined by Kaplan-Meier analysis (*P* = 0.0036; Fig. 4B). Notably, this survival benefit was achieved without measurable body weight loss, suggesting a favorable safety profile (fig. S3A). Treatment was initiated based on bioluminescence imaging (fig. S3B); however, tumor progression was subsequently monitored by abdominal palpation, as KPC cells are known to rapidly lose luciferase expression (20) due to their high proliferative capacity. This loss of luciferase activity does not compromise KPC-cell tumorigenicity or aggressiveness.

**Fig. 4.**
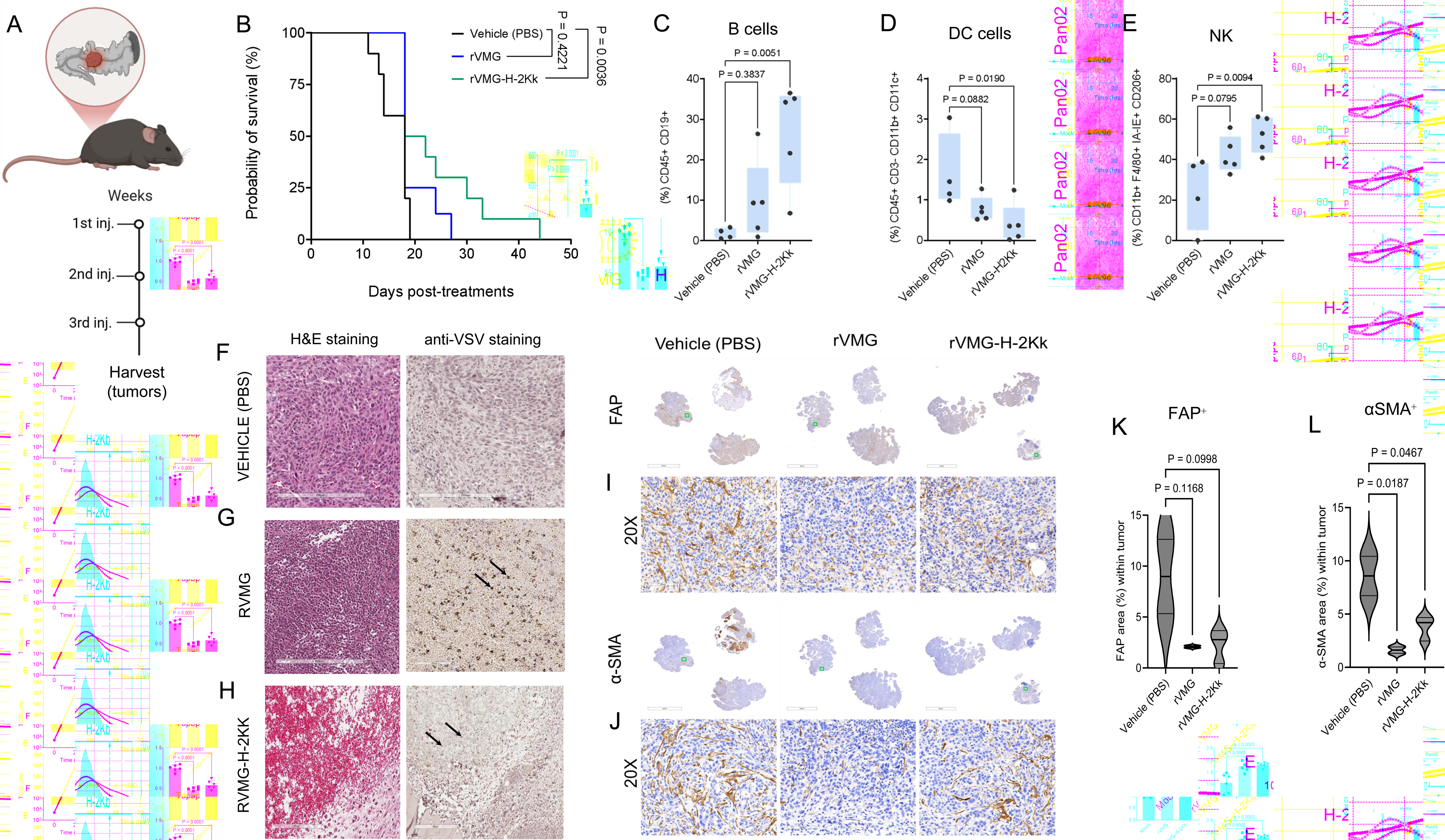
Treatment of an orthotopic metastatic KPC model with rVMG-H-2Kk. (**A**) Schematic of the treatment schedule for surgically implanted orthotopic KPC tumors. Mice received three intraperitoneal injections (100 µL each) of vehicle (PBS), rVMG, or rVMG-H-2Kk (1 x 10^7^ TCID_50_), and tumors were harvested 3 days after the final injection for downstream analyses. (**B**) Kaplan–Meier survival curves from an independent experiment comparing the 3 treatment groups. (**C** to **E**) Flow cytometry quantification of B cells, dendritic cells (DCs), and natural killer (NK) cells within the tumor microenvironment. (**F** to **H**) Representative H&E (left) and VSV-antigen immunohistochemistry (IHC; right) staining in tumor sections. Arrows represent positive areas for VSV antigens. (**I** and **J**) IHC staining for fibrosis markers: fibroblast activation protein (FAP; **I**) and α-smooth muscle actin (αSMA; j). (**K** and **L**) Quantification of FAP^+^ and αSMA^+^ cells in tumor sections. Bars represent mean ± SEM from three tumors per group. Statistical significance was determined by one-way ANOVA with *P* < 0.05 considered significant.

Intraperitoneal administration of rVMG-H-2Kk resulted in broad immunomodulation of the TME characterized by enhanced immune cell infiltration, reduced stromal fibrosis, and induction of antiviral and immunoregulatory pathways (list of antibodies in table S2). In particular, immunophenotypic analyses showed that rVMG-H-2Kk significantly increased the infiltration of B cells (*P* = 0.0051; Fig. 4C), reduced DC populations (*P* = 0.0190; Fig. 4D), and enhanced NK cells abundance (*P* = 0.0094; Fig. 4E) in the TME. Histopathologic evaluation and IHC demonstrated extensive necrosis and robust VSV-antigen positivity in tumors from both rVMG and rVMG-H-2Kk–treated mice (Fig. 4, F to H). Furthermore, treatment with rVMG vectors significantly reduced fibrosis as evidenced by decreased α-smooth muscle actin levels (rVMG, *P* = 0.0187; rVMG-H-2Kk, *P* = 0.0467; Fig. 4, J and L), while FAP levels tended to be lower (rVMG, *P* = 0.1168; rVMG-H-2Kk, *P* = 0.0998; Fig. 4, I and K). Further flow cytometric analysis of T cell activation and exhaustion markers demonstrated a reduced frequency of CD8 CD44 subset and CD4 /CD8, CD8 PD-1 /CD8, CD4 CD44 /CD8, and CD8 GzmB /CD8 T cell ratios in rVMG-H-2Kk–treated tumors (fig. S4A–E). Of particular interest, rVMG-H-2Kk reduced the proportion of PD-1 cells within the CD8 T cell, as reflected by the CD8^+^PD 1 /CD8 ratio, from ∼3.7% ±1.6% to undetectable levels (*P =* 0.0133; fig. S4C). Additionally, rVMG-H-2Kk significantly increased the frequency of CD45 CD3 cells (*P =* 0.0339; fig. S5A) within the TME. Analysis of CTL activation (ICOS^+^, CD44^+^ CD69^+^, and CD44^+^ Ki-67^+^) and immunosuppression (PD-1^+^, TIM-3^+^, LAG-3^+^, and CTLA-4^+^) markers in tumor infiltrates (fig. S6, A and B, respectively) revealed no significant differences among PBS-, rVMG-, and rVMG-H-2Kk–treated groups (all *P* > 0.05). In contrast, qPCR analyses showed upregulation of PDL-2 (*P* = 0.0146; fig. S7C) and important induction of genes involved in antiviral responses and T-cell recruitment (Mx1, IFNα, and IFNβ; all *P* < 0.0001; fig. S8, A to C) (list of primers in table S3). No statistically significant differences in hyaluronic acid or collagen content were observed, as assessed by Alcian blue^+^ (fig. S9, A and D) and picrosirius red^+^ staining (fig. S9, B and C).

### Systemic administration of rVMG-H-2Kk shows no evidence of toxicity in treated mice

To evaluate the safety of rVMG-H-2Kk, blood collected 3 days after the final treatment was analyzed, and toxicity data were compared with previously published normal mouse serum values.[30] We measured serum levels of key toxicity markers, including alanine transaminase, alkaline phosphatase, albumin, serum amylase, total bilirubin, blood urea nitrogen, total protein, creatinine, sodium, and phosphorus (fig. S10, A to J). Serum levels of potassium and calcium were below the limit of detection. No significant increases in serum toxicity markers were observed in either the rVMG or rVMG-H-2Kk groups compared to vehicle (PBS) controls (*P* > 0.05), indicating that rVMG-H-2Kk treatment is not associated with liver or kidney toxicity and supports its favorable safety profile in in vivo murine models. Specifically, levels of alanine aminotransferase and alkaline phosphatase, indicators of hepatocellular injury, as well as blood urea nitrogen and creatinine, markers of renal function, remained within normal ranges (fig. S10, A to I). Notably, a modest elevation in serum phosphorus was observed in the rVMG-H-2Kk group (fig. S10J), which may reflect increased tumor cell turnover or immune-mediated tumor lysis (21) rather than organ dysfunction and warrants further investigation.

### Spatial transcriptomic profiling reveals compartment-specific remodeling of the tumor microenvironment following rVMG-H-2Kk therapy in KPC tumors

To further characterize the effects of rVMG-H-2Kk therapy on the TME, we performed spatial transcriptomic profiling of orthotopic KPC tumors (three per group: vehicle [PBS], rVMG, and rVMG-H-2Kk) using the GeoMx Digital Spatial Profiler platform (Fig. 5A).

**Fig. 5.**
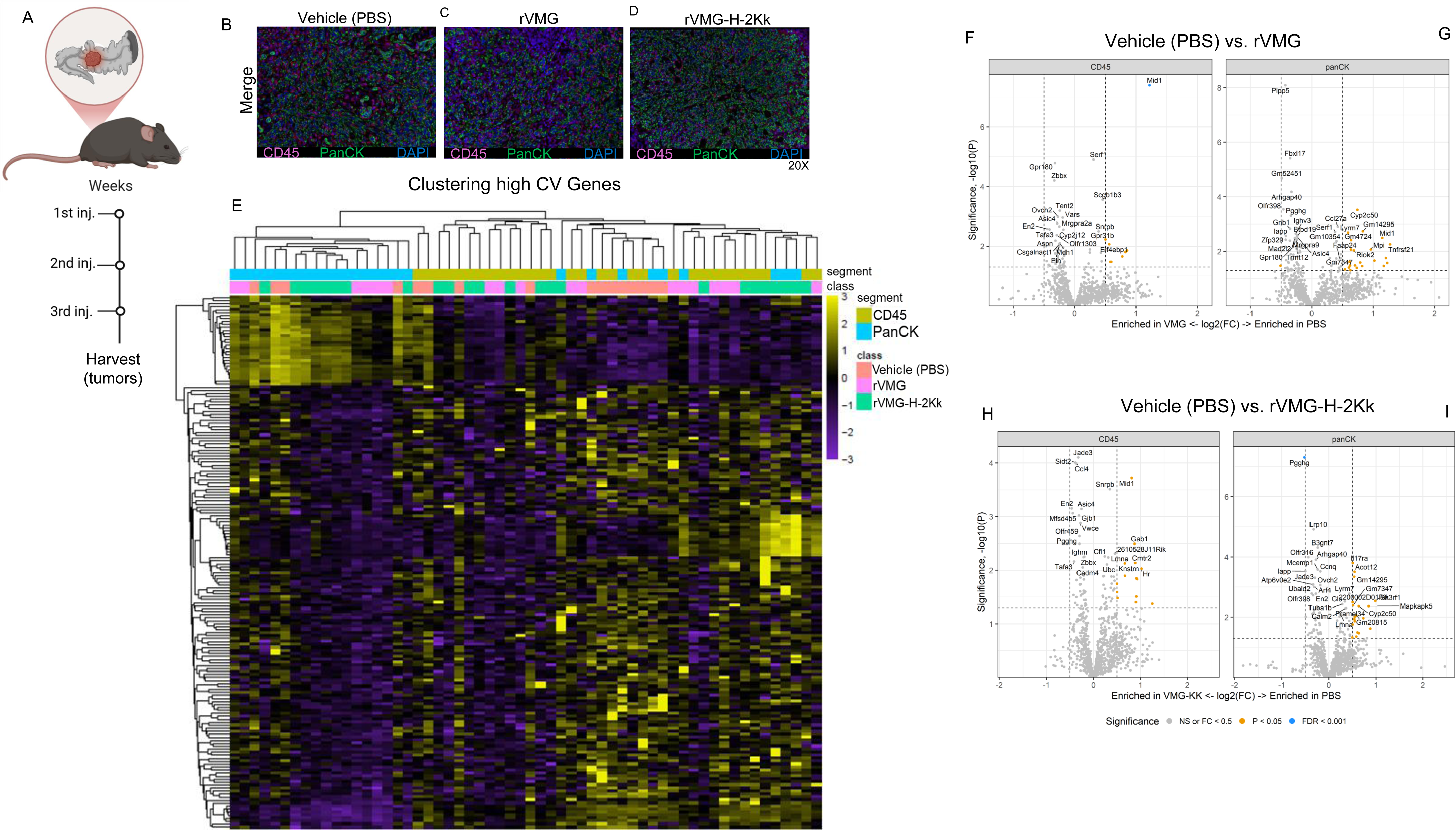
Spatial transcriptomic profiling of treated KPC tumors by GeoMx Digital Spatial Profiler. (**A**) Schematic of the treatment regimen in the orthotopic KPC model showing the injection schedule. (**B** to **D**) Representative immunofluorescence images of KPC tumor sections stained for CD45 and panCK. Regions of interest (ROIs) were selected and segmented into areas of illumination (AOIs) to distinguish immune (CD45^+^) from epithelial (panCK^+^) compartments across treatment groups (vehicle [PBS], rVMG, and rVMG-H-2Kk). (**E**) Unsupervised hierarchical clustering heatmap of genes with high variability, determined using the coefficient of variation (CV; SD/mean), highlighting transcripts that differ markedly across AOIs. (**F** to **H**) Volcano plots comparing differential gene expression between the vehicle (PBS) and rVMG (**F**) and between the vehicle (PBS) and rVMG-H-2Kk (**G**), with an integrated analysis presented in (**H**). Statistically significant genes (*P* < 0.05) are indicated.

Immunofluorescence-guided segmentation allowed precise delineation of immune (CD45) and epithelial (panCK) compartments (Fig. 5, B to D).

Dimensionality reduction using uniform manifold approximation and projection (UMAP) and t-distributed stochastic neighbor embedding (tSNE) revealed distinct treatment-associated transcriptomic profiles in each group (fig. S11, A and B). Unsupervised clustering of highly variable genes confirmed distinct expression profiles within immune and epithelial compartments (Fig. 5E). In the CD45 compartment, PBS- and rVMG-treated tumors showed limited transcriptional changes, with modest enrichment of genes such as *Eif4ebp1*, *Gpr31b*, and *Mid1* in the PBS group (P < 0.05; Fig. 5F). In comparison, rVMG-H-2Kk–treated tumors exhibited a slightly broader shift, including enrichment of genes associated with innate immune activity and T cell recruitment, such as *Mid1* and *Gab1* (P < 0.05; Fig. 5H). Mid1, a microtubule-associated E3 ubiquitin ligase, has been implicated in immune cell viability and microtubule stabilization, contributing to T-cell function and trafficking (22). Gab1 acts as a docking protein in cytokine and toll-like receptor signaling pathways and is essential for TLR3/4- and RIG-I–mediated antiviral responses (23, 24). These changes were accompanied by the enrichment of biological pathways related to antigen processing, leukocyte proliferation, cytokine signaling, and cell-cell adhesion (fig. S12A), supporting a shift toward an immunostimulatory microenvironment. When comparing PBS- and rVMG-treated tumors in the epithelial (panCK) compartment, we found that rVMG induced a distinct transcriptional profile. Enriched transcripts in PBS-treated tumors included *Tnfrsf21, Mid1*, and *Lymr7* (P < 0.05; Fig. 5G). Similarly, in the panCK compartment, comparison of PBS- and rVMG-H-2Kk–treated tumors revealed enrichment of transcripts such as *IL17ra*, a receptor for IL-17 family cytokines involved in mucosal immunity and tumor immune surveillance, and *Acot12* (P < 0.05; Fig. 5I), a peroxisomal thioesterase that modulates lipid metabolism and may suppress tumor-promoting metabolic programs (25–27).

To interpret high-plex transcriptomic data, we employed common dimension reduction techniques, including UMAP and t-SNE, non-orthogonally constrained projection methods that cluster samples based on overall gene expression patterns (fig. S13A, B). Using these methods alongside the analysis described above, we compared transcriptomic profiles between rVMG- and rVMG-H-2Kk–treated tumors. No enriched transcripts were observed in the CD45 compartment (fig. S13C); however, in the panCK compartment, enriched transcripts included *Erdr1*, *Olfr937,* and *Cbx8* in the rVMG-H-2Kk group (fig. S13D). *Erdr1* has been associated with cell migration and cancer metastasis, while *Cbx8*, a component of the polycomb repressive complex, is known to promote epithelial-to-mesenchymal transition and tumor progression (28, 29) These alterations suggest a reduction in immunoevasive and pro-metastatic signaling.

Gene ontology (GO) analysis comparing PBS vs. rVMG, PBS vs. rVMG-H-2Kk, and rVMG vs. rVMG-H-2Kk (fig. S12–S14) revealed significant enrichment of pathways involved in cytoplasmic translation, metabolic regulation, and suppression of apoptotic signaling (fig. S12B). These findings suggest that, compared to rVMG and PBS, rVMG-H-2Kk not only enhances immune accessibility but also reprograms epithelial metabolism to support antigen presentation and cellular survival. The induction of Mid1- and Gab1-driven antiviral and cytokine signaling, alongside repression of genes associated with immune evasion and metastasis, highlights a dual mechanism by which rVMG-H-2Kk enhances immune engagement and tumor clearance.

### Combination therapy with rVMG-H-2Kk and dual checkpoint blockade enhances survival and establishes durable immunity

Given the observed upregulation of immunosuppressive markers within the TME (fig. S7, A to E), we evaluated whether combinatorial immune checkpoint blockade (ICI) could mitigate immune evasion. Orthotopic KPC tumor–bearing mice received systemic administration of rVMG-H-2Kk, anti–PD-1, and anti–CTLA-4 antibodies, either as single agents or in combination. Survival and tumor rejection were assessed across treatment arms (N = 11-12 mice per group) to determine therapeutic efficacy. Kaplan-Meier survival analysis demonstrated significant differences among the treatment groups (vehicle [PBS], rVMG-H-2Kk, ICI [anti-PD-1 + anti-CTLA-4], and rVMG-H-2Kk + ICI [Combo]). Combination therapy markedly improved survival (*P* = 0.0007) compared to the vehicle control (Fig. 6A). By day 90, 27% of mice in the Combo group remained alive, whereas no long-term survivors were observed in the vehicle, rVMG-H-2Kk, or ICI monotherapy groups (fig. S15A, table S4). Importantly, this survival benefit was achieved without weight loss in the combination-treated mice (fig. S15B).

**Fig. 6.**
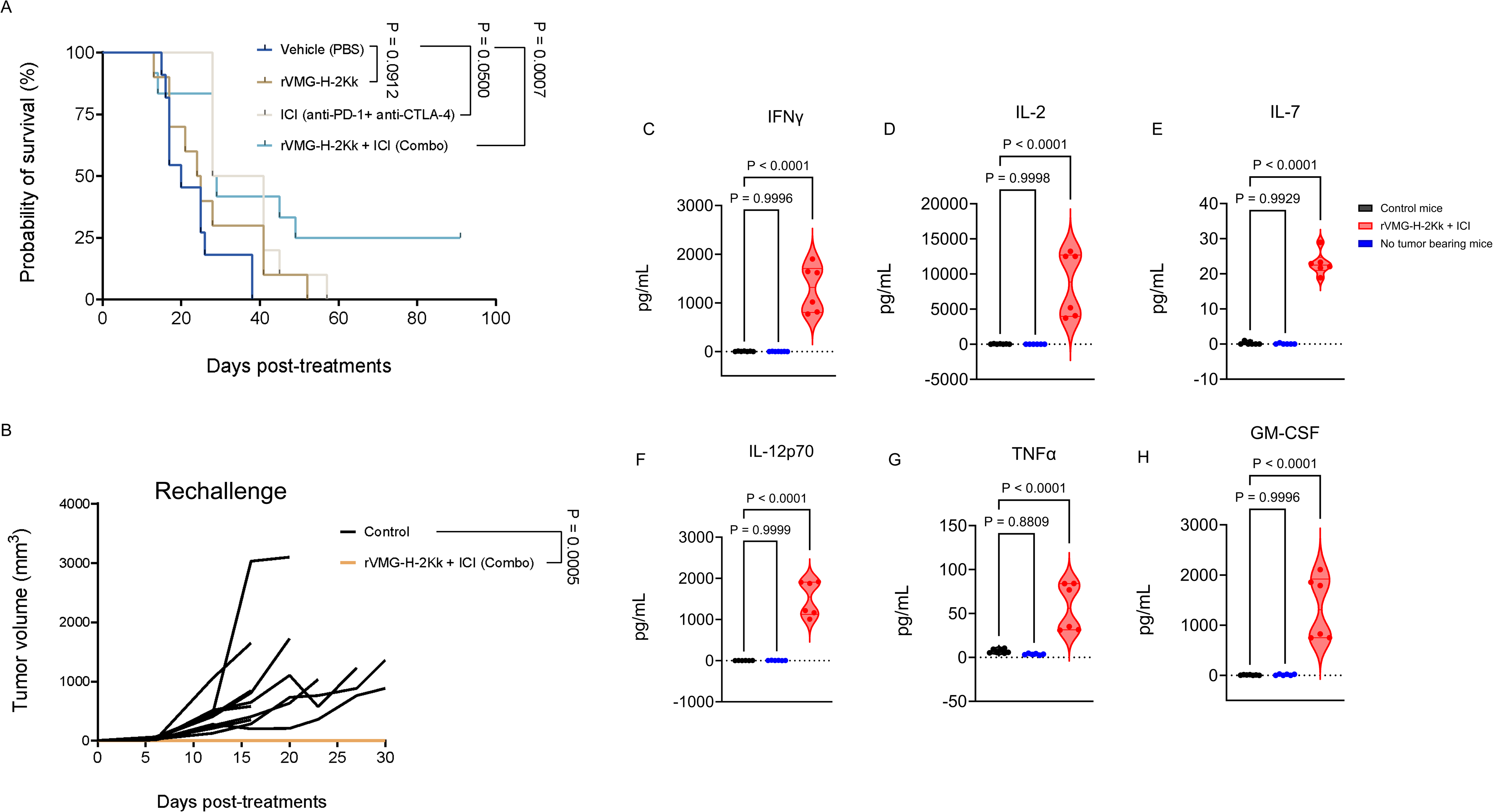
Combination of rVMGDHD2Kk with dual checkpoint blockade drives superior survival in an orthotopic KPC model. (**A**) Kaplan-Meier survival curves of mice treated with vehicle (PBS), rVMG-H-2Kk, ICI (anti-PD-1 + anti-CTLA-4), or rVMG-H-2Kk + ICI (Combo). Significant differences in survival were observed between groups (Log-rank test, *P* < 0.05). (**B)** Complete responders from the rVMG-H-2Kk + ICI (Combo) group were rechallenged subcutaneously with 2 × 10 KPC cells alongside ten age-matched naïve controls. Violin plots show serum levels of immune-related cytokines in long-term surviving mice treated with rVMG-H-2Kk + ICI (Combo, red), tumor-bearing untreated controls (black), and non–tumor-bearing controls (blue). Cytokines measured include **(C)** IFN-γ, **(D)** IL-2, **(E)** IL-7, **(F)** IL-12(p70), **(G)** TNF-α, and **(H)** GM-CSF, key mediators of T-cell activation, memory maintenance, and cytotoxic function. All rechallenged survivors remained tumor-free through day 30, whereas all naïve controls developed tumors by day 10. Serum samples were collected on day 30 post-rechallenge. Cytokine quantification was performed using the MILLIPLEX^®^ Mouse High Sensitivity T Cell Magnetic Bead Panel (MilliporeSigma) via the Luminex^®^ 200™ system. Data were log-transformed, normalized, and visualized in GraphPad Prism. Statistical significance was assessed by one-way ANOVA.

Building upon the improved survival observed with combination therapy, we next assessed whether surviving mice had developed durable, tumor-specific immune memory. Complete responders from the rVMG-H-2Kk + ICI (Combo) group were subjected to subcutaneous tumor rechallenge with 2 × 10 KPC cells alongside ten age-matched naïve controls. Strikingly, all three rechallenged mice from the Combo group resisted tumor growth through day 30, whereas all naïve controls developed palpable tumors by day 10 (Fig. 6B). These findings demonstrate that combination therapy not only prolongs survival but also induces durable antitumor immunity capable of mediating long-term protection against tumor recurrence.

### Systemic cytokine profiling identifies immune memory-associated proinflammatory remodeling in long-term survivors of rVMG-H-2Kk + ICI therapy

To delineate systemic immune correlates of durable tumor control, we performed multiplex cytokine profiling on serum collected from mice (N = 6) treated with rVMG-H-2Kk in combination with anti–PD-1 and anti–CTLA-4 that survived more than 90 days and resisted tumor rechallenge over an additional 30-day period. Sera from tumor-bearing untreated mice (N = 5) and non-tumor-bearing controls (N=5) were included for comparison. Quantitative analysis using Luminex xMAP^®^ technology revealed a distinct cytokine signature in combination-treated survivors, characterized by sustained elevation of proinflammatory and memory-supportive cytokines (Fig. 6, C to H; fig. S16, A to H; fig. S17, A to D). Notably, levels of IFN-γ, IL-2, IL-7, IL-12(p70), TNF-α, and GM-CSF were significantly increased in the rVMG-H-2Kk + ICI group relative to both control cohorts (*P* < 0.0001 for all; Fig. 6, C to H). These cytokines are well-established mediators of CD8 T cell survival, cytotoxic function, and memory differentiation.(30–32) IL-2 and IL-7 are essential for homeostatic maintenance of T cells(33, 34), IL-12(p70) drives Th1 polarization (35), and GM-CSF (36) promotes the differentiation and activation of myeloid cell populations. Additionally, we observed a robust upregulation of inflammatory mediators, including IL-1α, IL-1β, IL-6, CXCL1, CXCL2, CXCL5, CCL2, and IL-4 (*P* < 0.0001; fig. S16, A to H), indicating a proinflammatory state that may enhance immune cell recruitment and support effector T-cell activation and memory persistence. Analysis of regulatory and Th2-associated cytokines revealed elevated levels of IL-5, IL-10, IL-13, and IL-17A in survivors (*P* < 0.0001; fig. S17, A to D), suggesting a balanced immune activation profile. IL-10 has been implicated in the maintenance of memory T cell responses, regulatory T cell function, and modulation of inflammatory signaling.(37) Collectively, these data indicate that rVMG-H-2Kk + ICI therapy induces a robust systemic immune signature enriched for memory-supportive, inflammatory, and regulatory cytokines, correlating with complete tumor rejection upon rechallenge and supporting the generation of durable, functional antitumor immunity (Fig. 6B).

## DISCUSSION

In this study, we engineered an oncolytic vesiculovirus platform (rVMG-H-2Kk) to ectopically express the murine alloantigen H-2Kk, with the goal of inducing and redirecting host alloimmune responses toward tumor cells, thereby promoting immune-mediated tumor rejection. We evaluated this strategy using both locally delivered vectors in a subcutaneous model and systemically delivered vectors in an orthotopic PDAC model. In vitro, rVMG-H-2Kk retained robust replication and cytolytic function, driving strong upregulation of the ectopic transgene H-2Kk, endogenous H-2Kb [23, 29], and major components of the antigen presentation machinery, including β2 microglobulin, Tap1, and Tapbp. In vivo, rVMG-H-2Kk delayed tumor growth and extended survival in two independent syngeneic PDAC models.

To characterize the immunologic mechanisms underlying this response, we first examined IT delivery of rVMG-H-2Kk in the subcutaneous Pan02 model. We found that IT treatment with rVMG-H-2Kk delayed tumor progression and prolonged survival compared to parental rVMG and vehicle controls, suggesting that alloantigen delivery may amplify tumor-specific immune responses. Multiplex immunohistochemistry and flow cytometry analyses revealed a shift toward an immunostimulatory TME in both rVMG and rVMG-H-2Kk-treated tumors, marked by decreased FoxP3 regulatory T cells and increased granzyme B effector T cells.(38–40) These findings align with our previous reports that rVMG enhances CTL infiltration in murine PDAC TME, supporting its utility as a backbone for engineering rVMG-H-2Kk. (18, 19) However, the concurrent expansion of PD-1 exhausted CD8 T cells following rVMG-H-2Kk treatment pointed to an emerging compensatory shift toward immunosuppression within the TME.

Building on these findings, we evaluated systemic delivery of rVMG-H-2Kk in an orthotopic KPC model (41), which more closely recapitulates the metastatic and immune-excluded features of human PDAC. In this setting, rVMG-H-2Kk similarly extended survival and increased the infiltration of B cells, NK cells and a reduction in dendritic cells (DCs) populations. These changes are consistent with prior reports demonstrating that systemic administration of oncolytic viruses can inflame otherwise immune-cold pancreatic tumors and trigger tumor eradication.(4, 42) While B cells are often immunosuppressive in PDAC, their expansion here may reflect the formation of tertiary lymphoid structures or antigen-presenting subsets.(43, 44) The observed reduction in DCs could be due to viral targeting or migration(45), warranting further investigation. The increase in NK cells could suggest enhanced cytotoxic activity, further supporting the immunostimulatory effect of rVMG-H-2Kk.

To assess treatment-associated changes beyond immune infiltration, we conducted histopathologic and molecular analyses. KPC tumors from treated mice showed reduced stromal density, potentially facilitating CTL infiltration, while serum chemistry profiles confirmed no significant hepatotoxicity or nephrotoxicity. Molecularly, qPCR analyses revealed upregulation of *PD-L2* and type I interferon-responsive genes (*IFN*α*, IFN*β), which could indicate concurrent activation of antiviral and immunosuppressive pathways. Type I interferons are known to enhance antigen presentation while also inducing PD-L2 expression via STAT and IRF1 signaling (46), highlighting the dual role of rVMG-H-2Kk in promoting immunogenicity and eliciting adaptive immune resistance.

To further elucidate the mechanistic basis of rVMG-H-2Kk-induced remodeling, we performed spatial transcriptomic profiling of KPC tumors. Compartmental analysis of CD45^+^ immune and panCK^+^ epithelial cells revealed distinct transcriptional programs across treatment groups. When comparing PBS- and rVMG- or rVMG-H-2Kk–treated tumors, most of the enriched transcripts were observed in the PBS group. These include *Mid1*, a microtubule-associated E3 ubiquitin ligase, has been linked to immune cell viability and cytoskeletal stabilization (22), and *Gab1,* an adaptor gene that promote cell survival and participates in TLR3/4 and RIG-I-associated innate immune signaling.(23, 24)

Concurrently, transcriptional changes induced by rVMG-H-2Kk in the panCK compartment suggest a shift toward reduced immunoevasion and metastatic potential. Notably, enrichment of *IL17ra*, a receptor involved in IL-17–mediated mucosal immunity and tumor immune surveillance, points to enhanced immune activation within the TME. Upregulation of *Acot12,* a peroxisomal thioesterase that modulates lipid metabolism, implies metabolic reprogramming that may suppress tumor-promoting lipid pathways. (25–27) Additionally, enrichment of *Erdr1* and *Cbx8* (28, 29), genes associated with cell migration, epithelial-to-mesenchymal transition, and tumor progression, may reflect context-dependent immune remodeling or compensatory changes. The presence of *Olfr937*, though functionally uncharacterized in cancer, further highlights the distinct transcriptomic landscape induced by rVMG-H-2Kk. Gene ontology analysis revealed activation of cytoplasmic translation and metabolic regulation pathways alongside suppression of apoptotic programs, reflecting a coordinated, compartment-specific reprogramming of both immune and epithelial niches by rVMG and rVMG-H-2Kk.

Immune checkpoint inhibitors (ICIs) targeting PD-1/PDL-1 and CTLA-4 pathways have demonstrated synergistic activity across multiple solid tumors, including melanoma, renal cell carcinoma, and hepatocellular carcinoma, leading to FDA-approved combination regimens. (47–50) Since PD-1 interacts with both PD-L1 and PD-L2 ligands (51), we hypothesized that dual checkpoint blockade with anti-PD-1 and anti-CTLA-4 antibodies could counteract the emerging immunosuppressive axis observed in our in vivo models. Consistent with this, systemic administration of rVMG-H-2Kk combined with murine anti-PD-1 and anti-CTLA-4 antibodies significantly extended survival in the orthotopic KPC model and achieved complete tumor eradication in 27% of treated mice, compared to rVMG-H-2Kk or ICI monotherapy alone.

Importantly, all complete responders rejected subsequent tumor rechallenge, indicating the induction of durable and functional antitumor immune memory. Cytokine profiling of blood form the complete responders revealed a distinct immune signature enriched in memory- supportive and proinflammatory cytokines, including IFN-γ, IL-2, IL-7, IL-12(p70), GM-CSF, and TNF-α, which are critical for CD8 T cell persistence and recall responses.[38-40]

Elevated levels of IL-1α, IL-1β, IL-6, CXCL1, CXCL2, and CCL2 indicated a sustained immunostimulatory environment. In parallel, increased IL-10, IL-13, and may reflect regulatory tuning and tissue remodeling following tumor clearance. IL-10, in particular, has been associated with non-exhausted memory T-cell states and enhanced recall capacity. [44]

In summary, rVMG-H-2Kk virotherapy reprograms the immunosuppressive TME, enhances tumor immunogenicity, and synergizes with ICI to elicit durable antitumor immunity in murine PDAC. Ectopic alloantigen delivery provides a novel strategy to overcome neoantigen limitations and sensitize immune-excluded tumors. These findings support further clinical translation and refinement of this platform, including optimizing alloantigen selection, limiting antiviral clearance, and uncovering mechanisms of immune memory.

## MATERIALS AND METHODS

### Cell lines and culture conditions

Murine pancreatic ductal adenocarcinoma (KPC, Pan02), Vero, and baby hamster kidney (BHK- 21) cells were cultured in Dulbecco’s Modified Eagle’s Medium (DMEM) supplemented with 10% fetal bovine serum (Gibco), 1% penicillin/streptomycin (Gibco), and 2 mM L-glutamine (Gibco) in a humidified incubator with 5% CO_2_. Vero and BH-21 cells were obtained from the American Type Culture Collection, KPC cells were purchased from Kerafast, and Pan02 cells were obtained from the US National Cancer Institute (Frederick, MD, USA). Cells were routinely tested for mycoplasma contamination.

### Generation of recombinant oncolytic rVMG constructs

The oncolytic VSV-based vector rVMG was generated as previously described. [23] Recombinant constructs (rVMG), including vectors encoding murine MHC class I molecules, P04223-HA1K_MOUSE H-2Kk (rVMG-H-2Kk), were generated by inserting the respective transgene into the viral genome using a pVMG-XN2-based cloning strategy. All viral constructs were propagated in Vero or BHK-21 cells and purified using sucrose gradient ultracentrifugation. TCID_50_ values were determined by the Spearman-Kärber algorithm using serial viral dilutions in BHK-21 cells. Viruses were aliquoted and stored at −80°C. Aliquots were thawed immediately prior to use with less than five freeze-thaw cycles per aliquot.

### Impedance-based cell death analysis

KPC and Pan02 cells were seeded in E-Plate 16 VIEW plates at a density of 1 x 10^4^ cells per well and allowed to adhere overnight. Cells were infected at an MOI of 0.1 with rVMG or rVMG-H-2Kk in Gibco Opti-MEM and incubated for up to 72 hours at 37°C. Cell viability was assessed using the Agilent xCELLigence Real-Time Cell Analysis (RTCA) S16 device, which monitors real-time impedance-based cell proliferation and cytotoxicity. Data were analyzed using RTCA Software Pro (Agilent).

### One-step viral growth assay

To assess viral growth kinetics, 5 × 10^5^ Vero cells were seeded in 6-well plates containing 2 mL of complete DMEM. After overnight incubation, cells were mock infected or infected with rVMG or rVMG-H-2Kk (MOI 0.1) for 1 hour. Then, the inoculum was removed, cells were washed with PBS, and fresh medium was added. Supernatants were collected at 24 and 48 hours post-infection and stored at −80°C. Viral titers (TCID_50_) were determined by TCID_50_ values using serial viral dilutions in BHK-21 cells. Data are shown as the mean ± SEM from two independent experiments.

### Intratumoral injection of rVMG vectors in the Pan02 model

Murine Pan02 cells were maintained as described (see cell lines and culture conditions). For subcutaneous tumor implantation, 6- to 8-week-old male C57BL/6J (Strain #:000664) immunocompetent mice from Jackson Laboratories were injected in the right flank with 1 x 10^6^ Pan02 cells in 100 µL of sterile RPMI 1640 medium. Tumor volumes were monitored twice weekly using digital calipers, and volumes were calculated using the formula ½ x (length x width^2^). Once tumors reached a volume of approximately 80 to 120 mm^3^, mice (N = 10–12) were randomly assigned to treatment groups. Mice in the vehicle control group received 50 µL IT injections of sterile PBS once weekly for 3 consecutive weeks. Oncolytic rVMG groups received IT injections of 1 x 10^7^ TCID_50_ of rVMG or rVMG-H-2Kk in 50 µL PBS once per week for 3 weeks. The dose of 1 x 10^7^ TCID_50_ of rVMG was determined to be both safe and effective in our previous studies. [29] Tumor growth and body weight were recorded twice weekly. Three days after the final IT injection, mice designated for early endpoint analysis were humanely euthanized. Tumors (N = 3–5 per group) were excised for downstream analyses. For survival studies, mice were continuously monitored, and tumor and body weight were measured as previously described. Animals were observed for clinical signs and tumor burden until they reached humane endpoints or a tumor volume of 2,000 mm³ in accordance with institutional animal care and use regulations.

### Multiplex Immunohistochemistry

Pan02 tumors were fixed in 10% neutral-buffered formalin for 24 hours, then transferred to 70% ethanol for 72 hours, processed, and embedded in paraffin. Formalin-fixed paraffin-embedded (FFPE) tissue sections, 4 µm thick, were prepared and mounted onto positively charged glass slides. All IHC, including hematoxylin and eosin staining and multiplex IHC procedures, were performed by the UAMS Winthrop P. Rockefeller Cancer Institute Experimental Pathology Core using a Leica Bond RX automated stainer (Leica Microsystems). After deparaffinization and rehydration, antigen retrieval was conducted in either citrate buffer (pH 6.0) or EDTA buffer (pH 9.0), depending on the specific antibody requirements. Anti-VSV was purchased from Imanis Life Sciences (Cat#REA005) for anti-VSV IHC. A panel of primary antibodies was used to assess major immune markers, including CD8 (clone D4W2Z, Cell Signaling Technology, #98941), F4/80 (clone BM8, Cell Signaling Technology, #70076), Granzyme B (clone D6E9W, Cell Signaling Technology, #46890), MHC II (clone M5/114.15.2 or 14-5321-82, Invitrogen), FoxP3 (clone D6O8R, Cell Signaling Technology, #12653) (see Table 1). Each antibody staining was detected with an Opal fluorophore (Opal 480, 520, 570, 620, or 690; Akoya Biosciences), followed by an appropriate epitope retrieval cycle to strip residual immunoreagents while retaining the covalently bound fluorophore. DAPI (Sigma-Aldrich) served as a nuclear counterstain. Stained slides were mounted with SlowFade™ Gold antifade reagent (Thermo Fisher), and images were captured using the Akoya PhenoImager HT platform.

### IHC image analysis

Multiplex IHC image analysis was performed on three tumors per group using a HALO™ image analysis platform (Indica Labs). Large necrotic areas were excluded. The frequency of single-positive (CD8^+^ T cells, F4/80^+^ macrophages, MHCII^+^, and FoxP3^+^) and co-expressing cells (e.g., CD8^+^/F4/80^+^, CD8^+^/Granzyme B^+^) were quantified as cell counts per tissue area or as a percentage of total DAPI^+^ nuclei. Representative fields were selected for figures. Group comparisons were made using an unpaired Student’s t-test or one-way ANOVA when comparing more than two groups.

### Orthotopic implantation of KPC cells

Exponentially growing, bioluminescent mouse KPC tumor cells (derived from EUP012-FP; Kerafast) were harvested by trypsinization, washed in serum-free medium, and suspended in basement membrane–supplemented medium. The EUP012-FP cell line is derived from a mouse KPC tumor that incorporates common genetic mutations seen in human pancreatic cancer and demonstrates significant metastatic potential along with a highly fibrotic extracellular matrix.[42] Six- to eight-week-old male C57BL/6J mice (Strain #:000664, Jackson Laboratories) were anesthetized with 1.5 to 3% isoflurane, and the upper left quadrant was shaved and disinfected using alternating scrubs of betadine and alcohol. A 1-cm incision was made to expose the pancreas and spleen, allowing gentle externalization of the pancreas. Using a 27-gauge needle, 5 x 10^5^ EUP012-FP cells in 30 µL of medium were injected directly into the pancreatic parenchyma. A cotton-tipped applicator was briefly placed over the injection site to prevent leakage. The peritoneal wall and skin incisions were then closed with wound clips. For proper closure, wounds were inspected for hemostasis and debris, treated with betadine if necessary, and aligned before firmly squeezing the clip applicator. Removal was performed at an appropriate time point by aligning the removal tool at a ∼45° angle to open and extract the clips, confirming adequate healing. Postoperative analgesia was provided by the subcutaneous administration of Meloxicam (2 mg/kg) once or daily for up to 3 days. Mice were allowed to recover in a 37°C warming cage for at least 30 minutes prior to returning to clean home cages. All procedures were approved by the UAMS Institutional Animal Care and Use Committee (IACUC) and adhered to institutional policies on pain management and ethical endpoints.

### Statistical analysis

Data are presented as mean ± standard error of the mean (SEM). Statistical analysis was performed using GraphPad Prism (version 10.2.3; GraphPad Software, Inc). One-way ANOVA with Dunnett’s post hoc test was used for multiple comparisons among groups. Significance thresholds were set at *P* < 0.05. Survival differences among treatment groups were analyzed using the log-rank (Mantel-Cox) test. The Gehan-Breslow-Wilcoxon test, which gives greater weight to early events, supported early survival divergence between groups.

## Data and materials availability

All experimental protocols, raw data, and final reports have been archived and available to investigators upon request, ensuring full compliance with regulatory requirements.

## Acknowledgments

This work was supported by the National Institutes of Health (NIH)/National Cancer Institute (NCI) through a K01 award (CA234324 to B.M.N.), an American Association for Cancer Research (AACR) grant (to B.M.N.), a Seed of Science Award from the Winthrop P. Rockefeller Cancer Institute (to B.M.N.), and NIH DP2CA301099 (to B.M.N.). Additional support was provided by a Development Enhancement Awards for Proposals (DEAP) from the UAMS Research Committee (to B.M.N.) and the College of Medicine’s intramural grant Barton Pilot Award Program (to B.M.N.). This work was supported by funds through the NCI - Cancer Center Support Grant (CCSG) – P30CA134274. The funders had no role in study design, data collection, interpretation, or in the decision to submit the work for publication.

## Author contributions

A.C., M.Z.T., A.G.B., K.U.F., M.J.B., M.J.C., A.G.B., and B.M.N. contributed to the study concept and design, data acquisition, data analysis, data interpretation, and manuscript drafting. A.C., M.Z.T., Z.C., K.U.F., R.S.S., B.M., N.E.E., E.R., C.C.S., E.R.S., I.R.M., A.U., S.R.P., O.M., C.Y.C., R.G., V.Z.G., M.E.F.Z., M.J.B., M.J.C., A.G.B., M.A.B., and B.M.N. contributed to data acquisition, data analysis, data interpretation, drafting, and critical revision of the manuscript. M.A.B. and B.M.N. contributed to bioinformatic analysis. All authors approved the final, submitted version of the manuscript.

---

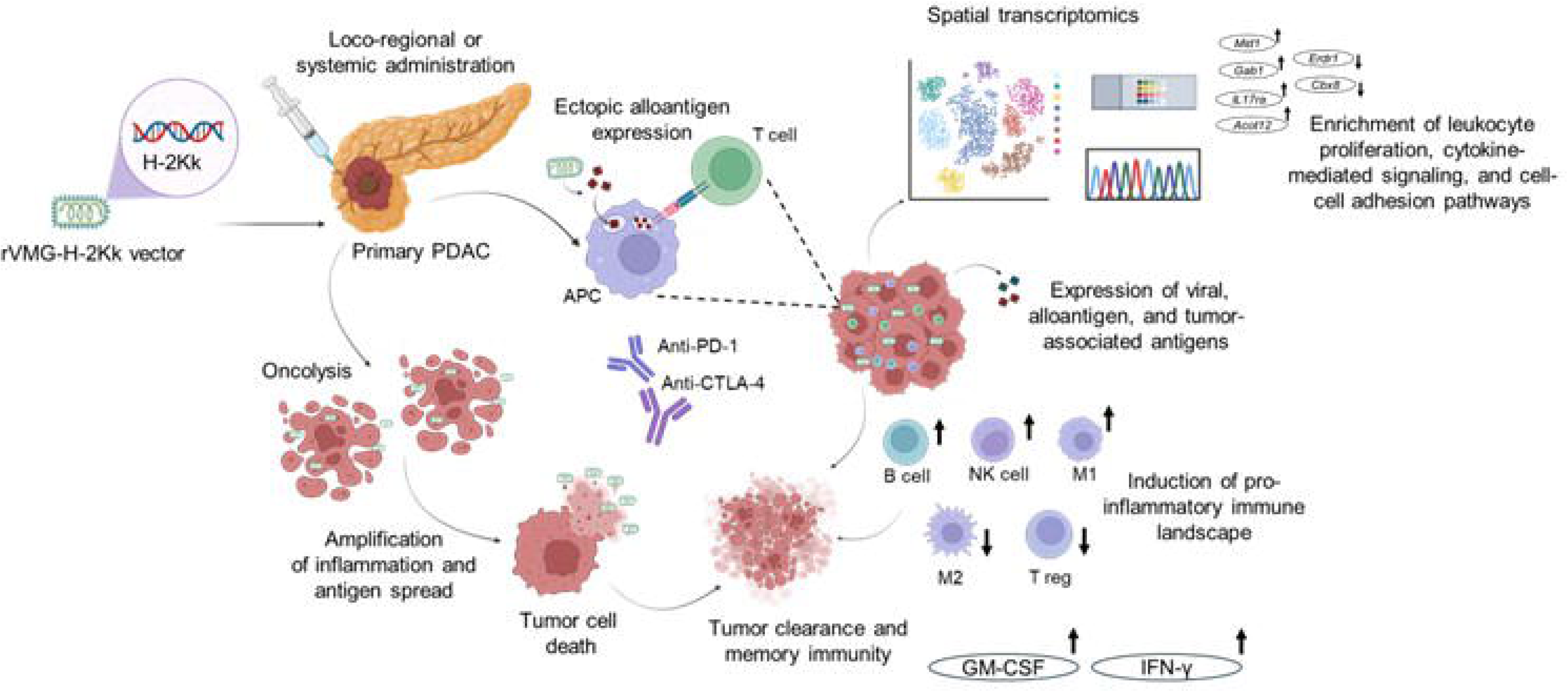

