## Supplemental methods and legends for "Pancreatic tumor microenvironment reprogramming via alloantigen-expressing virotherapy elicits tumor rejection and improves immunotherapy response"

Associate Professor Program in Oncology

Department of Pharmacology & Physiology, University of Maryland School of Medicine

Marlene and Stewart Greenebaum NCI Comprehensive Cancer Center, University of Maryland School of Medicine, 655 W. Baltimore St., Baltimore, MD, 21201, USA

Mitesh J. Borad, MD

Professor of Medicine

Division of Hematology and Medical Oncology, Mayo Clinic in Arizona

Mayo Clinic Comprehensive Cancer Center, Mayo Clinic, 5777 East Mayo Boulevard, Phoenix, AZ 85254, USA.

Text word count: 3,551

Supplementary figures:17

Supplementary table: 4

### SUPPLEMENTAL MATERIALS AND METHODS

#### Cell proliferation assay

KPC and Pan02 cell viability was assessed 72 hours post-infection with rVMG or rVMG-H-2Kk as previously described.[29] Cells were counted and evaluated for viability by trypan blue exclusion (Thermo Fisher) using a Luna-II Automated Cell Counter (Logos Biosystems). Triplicate cell samples were infected at an MOI of 10, 1, 0.1, 0.01, and 0 with rVMG or rVMG-H-2Kk in Gibco Opti-MEM and incubated for 72 hours at 37°C. At 72 hours, MTS reagent (CellTiter 96® AQueous One, Promega) was added and incubated for 1 hour. Absorbance was then measured at 490 nm on a Synergy HTX plate reader (BioTek) using Gen5 software (v3.12.08, Agilent Technologies).

#### Cytopathic effect assay

KPC cells were seeded in 6-well plates at a density of $5 \times 10^5$ cells per well, followed by infection at an MOI of 0.1. Cytopathic effects were monitored at 48 hours post-infection. Brightfield images were acquired using an EVOS XL Core microscope (Thermo Fisher Scientific). Infected cells were also fixed in 4% paraformaldehyde, stained with crystal violet (Sigma-Aldrich), and imaged to assess viral-induced cell lysis.

**SYBR green PCR analysis**

Total RNA was extracted from infected (MOI of 0.1 for 6 hours) and control KPC cells using the RNeasy Mini Kit (Qiagen) following the manufacturer's protocol. Synthesis of cDNA was performed using the iScript cDNA Synthesis Kit (Bio-Rad). SYBR Green-based real-time quantitative PCR (qPCR) was conducted using the Power SYBR Green PCR Master Mix (Thermo Fisher Scientific) on a QuantStudio 7 Flex Real-Time PCR System (Applied Biosystems). Gene expression levels were normalized to Rplp0 using mouse qPCR Primers and analyzed using the $\Delta\Delta$Ct method. Primer sequences for Rplp0 and all target genes are listed in table S3. Expression of H-2Kk and H-2Kb was measured using gene-specific primers, while additional targets included B2M, Tap1, Tapbp, VSV-N, MX1, IFN$\alpha$, IFN$\beta$, PD-1, PD-L1, CTLA-4, and LAG-3. All reactions were performed using validated mouse-specific primers to assess immune-related and viral transcripts relevant to treatment response. All experiments were repeated six times.

**Bioluminescence Imaging**

Mice were anesthetized with isoflurane and imaged once weekly using an IVIS Xenogen system to assess tumor growth and viral biodistribution. Isoflurane (2–5%) was first administered in an induction chamber; animals were then transferred to a heated imaging stage and fitted with a nose cone delivering 1.5–2% isoflurane to maintain an appropriate anesthetic plane throughout the procedure. Additional warming measures (e.g., heating pads and heat lamps) were employed as needed to preserve core body temperature. Before each imaging session, mice received an intraperitoneal injection of D-luciferin (150 mg/kg in ~10 µL/g body weight, dissolved in sterile water). Anesthetized mice were placed ventrally on the imaging platform, and bioluminescence signals were recorded for <10 minutes per cohort, minimizing stress and precluding the need for restraints. The respiratory rate was monitored continuously, and the isoflurane concentration was adjusted as necessary to maintain stable anesthesia. This non-invasive imaging approach allowed real-time tracking of virus localization without imposing additional physical burden on the animals.

**Tumor dissociation and flow cytometry**

Pan02 and KPC tumors were dissociated using a gentleMACS™ Dissociator (Miltenyi Biotec) with the Mouse Tumor Dissociation Kit (Miltenyi Biotec), following the manufacturer's protocol. The resulting cell suspensions were passed through a 30 µm cell strainer (Miltenyi Biotec), then washed in PBS with 1% fetal calf serum (FCS) (ThermoFisher). Cells were pelleted by centrifugation at 500 × g for 5 min at 25°C, resuspended in PBS + 1% FCS, and counted using trypan blue on an Invitrogen Countess 3 Automated Cell Counter (ThermoFisher). For immunostaining, cells were prepared at 1 x $10^6$ per tube and incubated with 0.5 µL of live/dead stain per 1 x $10^6$ and 1 µL of each fluorochrome-conjugated antibody (see Table 2 for detailed antibody panel) per 1 x $10^6$ cells in a total volume of ~100 µL for 30 min at 4°C in the dark. After staining, cells were washed twice with PBS under the same centrifugation conditions. The pellet was resuspended in 100 µL of PBS, and 5 µL of True-Stain Monocyte Blocker (Cat# 426102, BioLegend) was added to block non-specific binding of PE/Dazzle 594, PE/Cyanine7, and APC/Fire 750. Cells were then fixed (according to the respective fixative kit instructions) and immediately analyzed on a Cytek Northern Lights flow cytometer at the UAMS Flow

Cytometry Core. Flow Cytometry data were processed using FlowJo software (v10.10, BD Biosciences).

**Flow cytometry analysis**

Surface expression of MHC class I molecules was analyzed by flow cytometry at the University of Arkansas for Medical Sciences (UAMS) Flow cytometry Core. Infected and control KPC cells were detached using Accutase (Sigma-Aldrich), washed, and stained with fluorescently labeled anti-mouse H-2Kk Antibody clone 36-7-5 (BioLegend) or anti-mouse H-2Kb Antibody clone AF6-88.5 (BioLegend) for 30 minutes at 4°C. Cells were washed and then analyzed using a BD FACSCelesta flow cytometer (BD Biosciences). Data were analyzed using FlowJo software (v10.10, BD Biosciences).

**Survival analysis following intraperitoneal oncolytic virus administration**

Once orthotopic tumors reached approximately 3 mm in diameter on day 7 (verified by palpation or bioluminescence imaging [BLI]), mice (N = 10–12) were randomly assigned to receive intraperitoneal injections of vehicle (PBS), rVMG, or rVMG-H-2Kk at 1 x $10^7$ $TCID_{50}$ (diluted in 100 µL of PBS), once weekly for 3 weeks. Mice were imaged at designated intervals (e.g., daily for 1 week, then weekly) using BLI to assess tumor burden and therapeutic efficacy. Clinical signs (body weight, appearance, behavior) were monitored, with humane endpoints defined by institutional guidelines. At the study endpoint or when meeting humane criteria, mice were euthanized per IACUC-approved protocols, and tumors were collected for downstream assays.

**Integrated whole transcriptome profiling of FFPE KPC tumors using GeoMx digital spatial profiling and next-generation sequencing**

Whole transcriptome assay was performed on three separate FFPE KPC tumor samples, employing an integrated workflow that encompassed RNA slide preparation, library construction with rigorous quality control, next-generation sequencing, and comprehensive data analysis. FFPE blocks were sectioned at 4 µm, mounted on glass slides, and baked at 60°C for 60 minutes prior to processing on a BOND RX instrument. In situ hybridization was conducted using a 200 µL hybridization solution with overnight incubation at 37°C, followed by stringent washes, blocking, and morphology marker application. Regions of interest were precisely delineated on the GeoMx Digital Spatial Profiler, where targeted oligonucleotides were cleaved by UV light and collected for further processing. Subsequently, the collected products underwent PCR amplification with indexing, and libraries were purified using AMPure XP beads, quantified via Qubit 3.0, and size-verified on an Agilent BioAnalyzer, yielding an expected amplicon size of approximately 162 bp. Sequencing was carried out on a NovaSeq 6000 system with paired-end 2 x 150 bp reads following established Illumina protocols. Data analysis involved stringent quality control at both the segment and probe levels, including filtering based on raw read counts, alignment, trimming, and sequencing saturation. The limit of quantification was determined for each segment using the geometric mean and standard deviation of negative control probes, and normalization was achieved through the upper quartile method. Exploratory analyses, including dimensionality reduction (UMAP, tSNE), clustering of variable genes, and differential expression via linear mixed-effect models, were employed to rigorously interrogate the data.

**Bioinformatics analysis**

Spatial transcriptomic data were generated using the GeoMx Digital Spatial Profiler platform, enabling high-plex profiling of over 18,000 protein-coding genes from murine tissue sections. Using a whole mouse transcriptome atlas assay, tissue sections were stained for markers such as CD45 and panCK, and regions of interest were segmented into areas of illumination to differentiate immune from epithelial compartments across four treatment groups (vehicle [PBS], rVMG, and rVMG-H-2Kk). Quality control was conducted at both the segment and probe levels. Segments with <1,000 raw reads or with alignment, trimming, or stitching rates below 80% were excluded, and those with sequencing saturation below ~50% were flagged. Background signal was assessed via negative control and no-template control counts, while morphological criteria such as nuclei count (>100 per segment) and area were also considered. At the probe level, Grubb's test identified outliers; probes were removed if the geometric mean ratio of probe counts to overall target counts was <0.1 or if flagged in >20% of segments. Gene-level counts were computed as the geometric mean of multiple probes, and the limit of quantification was determined by using negative control probe metrics. Normalization was achieved with the upper quartile method to correct for technical variability, ensuring consistent expression distributions across segments. Unsupervised analyses, including UMAP, t-SNE, and hierarchical clustering of high variability genes (determined via coefficient of variation), were performed to explore spatial clustering and gene expression heterogeneity. Differential expression analysis employed linear mixed-effect models with treatment as fixed effects and tissue samples as random effects to compare groups (e.g., PBS vs. rVMG, PBS vs. rVMG-H-2Kk, and rVMG vs. rVMG-H-2Kk), and results were visualized using volcano plots. Finally, enrichment analysis was performed to elucidate the biological pathways underlying the observed differential expression. Gene ontology biological process terms were used as a reference, and enrichment was statistically evaluated using standard methods such as Fisher's exact test.

**Serum biochemistry analysis**

Blood chemistry analysis was performed by the UAMS DNA Damage and Toxicology Core using an Abaxis Piccolo Xpress chemistry analyzer (Abaxis) to evaluate liver toxicity markers (aspartate transaminase, alkaline phosphatase, albumin), nephrotoxicity markers (creatinine, blood urea nitrogen), and serum electrolytes.

**Combination of rVMG-H-2Kk and dual checkpoint inhibitors in an orthotopic KPC pancreatic cancer model**

Six- to eight-week-old C57BL/6 mice (Strain #:000664, Jackson Laboratories) were orthotopically implanted with 5 x $10^5$ bioluminescent KPC (EUP012-FP) cells into the pancreas as described above. When tumors reached ~3 mm in diameter on day 7, mice (N = 11–12 per group) were randomized to receive IP injections of vehicle (PBS, 100 µL); rVMG-H-2Kk

(1 x $10^7$ $TCID_{50}$ in 100 µL); dual ICIs—InVivoPlus anti-mouse PD-1 (CD279; clone 29F.1A12; Cat# BP0273) and InVivoPlus anti-mouse CTLA-4 (CD152; clone 9H10; Cat# BE0131)—each at 5 mg/kg in 100 µL (dual ICI)—or the combination of rVMG-H-2Kk plus both antibodies (Combo). Checkpoint inhibitors were administered twice weekly for 2 weeks, and the virus was given once weekly for 3 weeks. Animals were monitored biweekly for survival and body weight and were euthanized according to institutional guidelines. Complete responders (one dual ICI and three Combo group mice) were rechallenged on day 60 with $2 \times 10^6$ KPC cells subcutaneously, alongside 10 age-matched naïve controls. Tumor growth was measured twice weekly. Survival curves were compared by log-rank test, and longitudinal body weight and tumor volume data were analyzed by two-way ANOVA with Bonferroni correction. Statistical significance was defined as $P < 0.05$.

**Multiplex cytokine and chemokine profiling**

Quantitative profiling of murine cytokines, chemokines, and growth factors was performed using Luminex® xMAP® technology, allowing simultaneous detection of multiple analytes from limited serum volumes. All assays were conducted by the DNA Damage and Toxicology Core Facility at UAMS in collaboration with Eve Technologies Corporation. Serum samples were collected from three long-term surviving mice in the combination treatment group (rVMG-H-2Kk + immune checkpoint inhibitors) from the orthotopic KPC tumor study described in Fig. 6, as well as from five tumor-bearing control and five non–tumor-bearing control mice. Sample analysis was performed using the Luminex® 200™ system (Luminex Corporation) with Bio-Plex Manager™ software (Bio-Rad Laboratories). The Mouse High Sensitivity T-Cell 18-Plex

Discovery Assay® (MDHSTC18; Eve Technologies) was conducted according to the manufacturer's protocol (MILLIPLEX® Mouse High Sensitivity T Cell Magnetic Bead Panel, Cat# MHSTCMAG-70K, MilliporeSigma). The assay measured the following 18 analytes in 100µL of serum in triplicate: GM-CSF, IFN-γ, IL-1α, IL-1β, IL-2, IL-4, IL-5, IL-6, IL-7, IL-10, IL-12(p70), IL-13, IL-17A, CXCL1, LIX, CCL2, CXCL2, and TNF-α. Assay sensitivities ranged from 0.06 to 9.06 pg/mL, with analyte-specific detection limits provided in the manufacturer's documentation. Raw cytokine concentration data (pg/mL) were log-transformed and normalized across samples. Data visualization and statistical analyses were performed using GraphPad Prism (GraphPad Software). Violin plots were generated to compare cytokine profiles and identify immune correlates of rVMG-H-2Kk + ICI–mediated long-term tumor rejection. Statistical significance was assessed using one-way ANOVA with post hoc multiple comparisons or unpaired two-tailed t-tests, as appropriate. Statistical significance was defined as $P < 0.05$.

### SUPPLEMENTAL FIGURES AND TABLES

**Figure S1. Cell proliferation assay**. (**A**) KPC and (**B**) Pan02 cells were infected with rVMG or rVMG-H-2Kk at a multiplicity of infection of 10, 1, 0.1, or 0 (mock-infected) and incubated for 72 hours. Cell viability/proliferation was then assessed using an MTS assay (Promega). (**C**) Mice bearing tumors were treated with Vehicle (PBS), rVMG, or rVMG-H-2Kk (see Methods), and average body weight was recorded at the indicated time points. Data points in panel C represent mean values (N = 10–12 mice per group). Bars in panel A and B represent the mean ± SEM of triplicate wells.

**Figure S2. Gating strategy and tumor-infiltrating lymphocyte (TIL) phenotypes in subcutaneous Pan02 tumors.** (**A** and **B**) Representative gating strategy used to identify CD8$^+$ and CD4$^+$ T cells, as well as CD44$^+$ and PD-1$^+$ TILs in the tumor microenvironment. (**C** to **H**) Quantification of total T cells, CD4$^+$, CD8$^+$, CD8$^+$ GZmb$^+$, CD8$^+$ PD-1$^+$, and Tfh populations across different treatment groups. Boxes span the range of values from three tumors per group. Statistical significance was determined by one-way ANOVA with Dunnett's post-hoc with $P <$ 0.05 considered significant.

**Figure S3. Body weight monitoring in mice bearing KPC tumors.** (**A**) Mice bearing tumors (see Methods) were assigned to the indicated treatment groups and monitored for body weight over time. Each data point represents mean ± SEM (N = 10–12). Error bars denote SEM. (**B**) Examples of bioluminescent imaging over the course of the study.

**Figure S4. Analysis of immune cell subsets in KPC tumors.** (**A** to **E**) Bar graphs showing the frequency of indicated immune populations in tumor-bearing mice treated with Vehicle (PBS), rVMG, or rVMG-H-2Kk. Tumors were harvested, processed for flow cytometry, and gated as described (see Methods). Immune populations quantified included (**A**) CD8$^+$ CD44$^+$, (**B**)

CD4$^+$/CD8$^+$ ratio, (**C**) CD8$^+$ PD-1$^+$/CD8$^+$, (D) CD44$^+$ CD8$^+$/CD8$^+$ and (**E**) CD44$^+$ GzmB$^+$/CD8$^+$. Bars represent mean ± SEM from three tumors per group. Statistical significance was determined by one-way ANOVA with Dunnett's post-hoc with $P < 0.05$ considered significant.

**Figure S5. Flow cytometry analysis of immune populations in treated KPC tumors.** Tumors from mice treated with Vehicle (PBS), rVMG, or rVMG-H-2Kk were harvested at the indicated endpoint, and single-cell suspensions were analyzed by flow cytometry. Immune populations quantified included total T cells (CD45$^+$CD3$^+$, [**A**]) and (**B**) CD45$^+$CD3$^-$ cells, (**C**) CD8$^+$ T cells, (**D**) CD4$^+$ T cells, (**E**) NK cells, (**F**) macrophages, (**G**) M1 macrophages, (**H**) monocytes and (**I**) neutrophils. Data are presented as box plots overlaid with data points from 4 vehicle-group tumors (all panels), 5 rVMG-group tumors (all panels), and either 4 rVMG-H-2Kk tumors (panels A through C) or 5 rVMG-H-2Kk tumors (panels **D** through **I**). Statistical significance was determined using one-way ANOVA with Dunnett's post-hoc with $P < 0.05$ considered significant.

**Figure S6. T-cell activation and exhaustion marker profiling in treated KPC tumors.** (**A**) Graph showing the frequency of T-cell activation markers (ICOS, CD44, CD69 and Ki-67$^+$) in KPC tumors from mice treated with Vehicle (PBS), rVMG, rVMG-H-2Kk, or dual immune checkpoint blockade. (**B**) Graph showing the expression of T-cell exhaustion markers (PD-1, LAG3, and TIM3) across the same treatment groups. Tumor samples were collected at the specified endpoint and analyzed using a multiplex flow cytometry assay. Data are shown as box plots overlaid with data points from N= 4-5 tumors per group. Statistical significance was determined by one-way ANOVA ($P < 0.05$).

**Figure S7. qPCR analysis of immunosuppressive markers in KPC tumors.** Relative mRNA expression of major immunosuppressive markers was quantified by qPCR in KPC tumors from mice treated with Vehicle (PBS), rVMG, or rVMG-H-2Kk. The markers analyzed include (**A**) PD-1, (**B**) PDL-1, (**C**) PDL-2, (**D**) CTLA-4 and (**E**) LAG-3. Data are shown as fold change in log2 units relative to the mouse Rplp0 gene (N= 3 tumors per group). Statistical significance was determined by one-way ANOVA ($P < 0.05$).

**Figure S8. qPCR analysis of interferon-stimulated genes (ISGs) in KPC tumors**. (**A**) Relative expression of the ISG Mx1 in KPC tumors from mice treated with Vehicle (PBS), rVMG, or rVMG-H-2Kk. (**B** and **C**) qPCR analysis of antiviral interferon gene (IFNα and IFNβ) expression in the same tumor samples. Data are shown as fold change in log2 units relative to the mouse Rplp0 gene and represent (N = 3–6 tumors per group). Statistical significance was determined by one-way ANOVA ($P < 0.05$).

**Figure S9. Analysis of stromal components in KPC tumors.** (**A** and **B**) Quantification of two major stromal markers, hyaluronic acid (Alcian blue staining) and collagen (Picro Sirius Red staining[PSR]), in tumor sections from mice treated with Vehicle (PBS), rVMG, or rVMG-H-2Kk. Data are presented as violin plots showing the distribution of staining intensity or area across treatment groups. (**C** and **D**) Representative images of Alcian blue and Picro Sirius Red staining, respectively, showing hyaluronic acid and collagen deposition within the tumor microenvironment. Statistical significance was determined by one-way ANOVA ($P < 0.05$).

**Figure S10. Serum toxicity markers in orthotopic kpc mice following oncolytic therapy**. Orthotopic KPC tumor-bearing mice were treated as described in Figure 4, and blood was collected 3 days after the final treatment. Untreated mice served as controls. Panels show serum levels of key toxicity markers: (**A**) alanine transaminase (ALT), (**B**) alkaline phosphatase (ALP), (c) albumin , (**D**) serum amylase, (**E**) total bilirubin, (**F**) serum blood urea nitrogen (BUN), (**G**) total protein, (**H**) creatinine, (**I**) sodium, and (**J**) phosphorus. Serum levels of potassium ($K^+$) and calcium ($Ca^+$) were below the limit of detection (data not shown). Bars represent mean ± SEM from either four mice per group (vehicle (PBS) and no treatment) or five mice per group (rVMG and rVMG-H-2Kk). Statistical significance was determined by one-way ANOVA with Dunnett's procedure to compare non-Vehicle groups to the Vehicle group.

**Figure S11. Dimensionality reduction and differential expression analysis of high-plex transcriptomic data.** (**A** and **B**) uniform manifold approximation and projection (UMAP) and t-distributed stochastic neighbor embedding (t-SNE) plots depicting overall gene expression profiles from Vehicle (PBS), rVMG, and rVMG-H-2Kk treatment groups. Both UMAP and t-SNE are non-orthogonally constrained projections that cluster samples based on global transcriptomic patterns. (**C** and **D**) Volcano plots showing differentially expressed transcripts between rVMG- and rVMG-H-2Kk–treated samples. Significant genes ($P < 0.05$) are highlighted.

**Figure S12. Gene ontology biological process enrichment analysis.** Differentially expressed genes from Vehicle (PBS)- versus rVMG-H-2Kk–treated tumors were analyzed using the gene ontology biological process database. Ridgeline plots of the same gene ontology enrichment data display the distribution of adjusted *P* values across significantly enriched pathways, highlighting key biological processes modulated by rVMG-H-2Kk treatment in (**A**) $CD45^+$ segments and (**B**) $panCK^+$ segments. Enriched pathways with adjusted $P$ values < 0.05 are shown.

**Figure S13. Gene ontology biological process enrichment in CD45$^+$ and panCK$^+$ tumor segments.** (**A**) Ridgeline plots illustrating enriched pathways in CD45$^+$ tumor segments when comparing rVMG-H-2Kk– and PBS-treated samples. (**B**) Ridgeline plots of enriched pathways in panCK$^+$ segments under the same comparison. Color indicates the adjusted *P* value for each pathway.

**Figure S14. Gene ontology biological process enrichment in CD45$^+$ and panCK$^+$ tumor segments**. (**A**) Ridgeline plots showing significantly enriched gene ontology biological processes in CD45$^+$ segments from rVMG-H-2Kk–treated tumors compared to rVMG-treated controls. (**B**) Ridgeline plots illustrating enriched gene ontology processes in panCK$^+$ segments under the same comparison. Enrichment analysis was performed on differentially expressed genes, and color intensity reflects the adjusted *P* value for each pathway.

**Figure S15. Combination therapy prolongs survival without inducing toxicity**. (**A**) Bar graph showing the proportion of mice surviving beyond 90 days with no detectable tumor burden. (**B**) Longitudinal tracking of body weight across treatment cohorts. No group experienced ≥10% weight loss during the study period, indicating a favorable safety profile.

**Figure S16. Inflammatory cytokines elevated in rVMG-H-2Kk + ICI–treated survivors.** Violin plots showing serum levels of (**A**) IL-1α, (**B**) IL-1β, (**C**) CXCL2, (**D**) CXCL5, (**E**) IL-6, (**F**) CXCL1, (**G**) CCL2 and (**H**) IL-4 in sera from rVMG-H-2Kk + ICI-treated mice (Combo, red), tumor-bearing controls (black), and non–tumor-bearing controls (blue). Serum was collected on day 30 post-rechallenge. Assays were performed using the MILLIPLEX® panel on the Luminex® 200™ platform. Statistical significance was assessed by one-way ANOVA.

**Figure S17. Regulatory cytokines in rVMG-H-2Kk + ICI–treated survivors.** Violin plots showing serum levels of (**A**) IL-5, (**B**) IL-10, (**C**) IL-13, and (**D**) IL-17A mice treated with rVMG-H-2Kk + ICI (Combo, red), tumor-bearing controls (black), and non–tumor-bearing controls (blue). Serum was collected on day 30 post-rechallenge. Cytokine measurements were performed using the MILLIPLEX® panel on the Luminex® 200™ platform. Statistical significance was assessed by one-way ANOVA.

**Supplemental tables**

**Supplemental Table 1. List of antibodies (IHC).** The following antibodies were used for multiplex IHC: anti-CD8, anti-F4/80, anti-FoxP3, anti-MHC II, and anti-Granzyme B. Data are presented as mean ± SEM from three tumors per group.

**Supplemental Table 2. Antibodies used for flow cytometry analysis.** This table lists the antibodies employed for flow cytometric characterization of immune cell subsets in KPC tumor experiments, including target cell surface marker, clone, fluorochrome, and manufacturer. All staining was performed following the manufacturer's recommendations, with appropriate isotype controls to ensure specificity.

**Supplemental Table 3. Mouse qPCR primer sequences.** Forward and backward primer sequences are shown for the reference gene Rplp0 and 14 non-reference genes.

**Supplemental Table 4. Long-term survival defined as survival without detectable tumor burden beyond 90 days.** Summary of median survival times and percentage of long-term survivors (defined as survival beyond 90 days) across treatment groups in the orthotopic KPC murine model. No long-term survivors were observed in the Vehicle, rVMG-H-2Kk, or ICI groups, whereas combination therapy achieved 28% long-term survival.
