## Supplemental figures and tables for "Pancreatic tumor microenvironment reprogramming via alloantigen-expressing virotherapy elicits tumor rejection and improves immunotherapy response"

### Supplemental figures & tables

### Supplemental Fig. 1

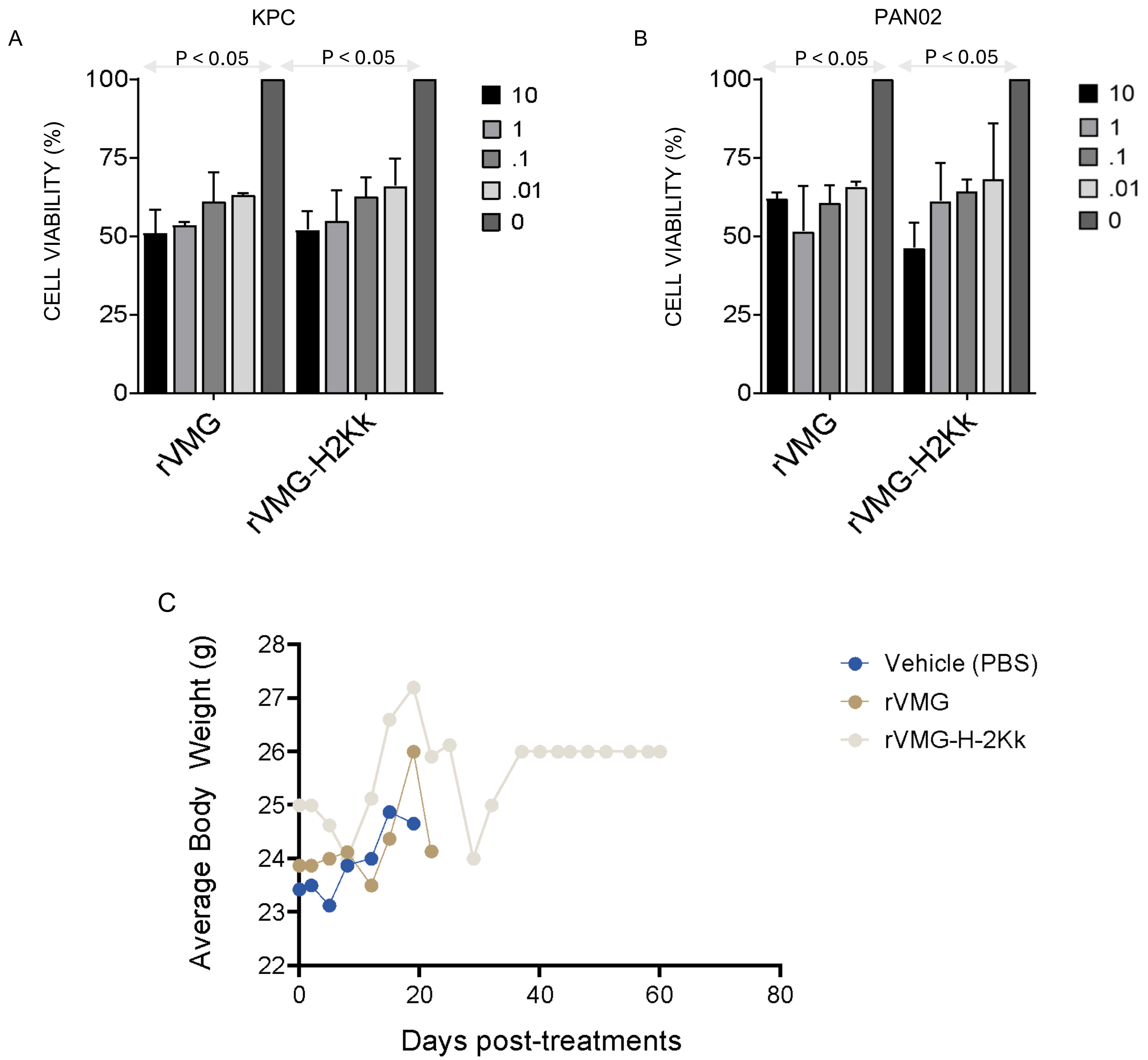

### Supplemental Fig. 2

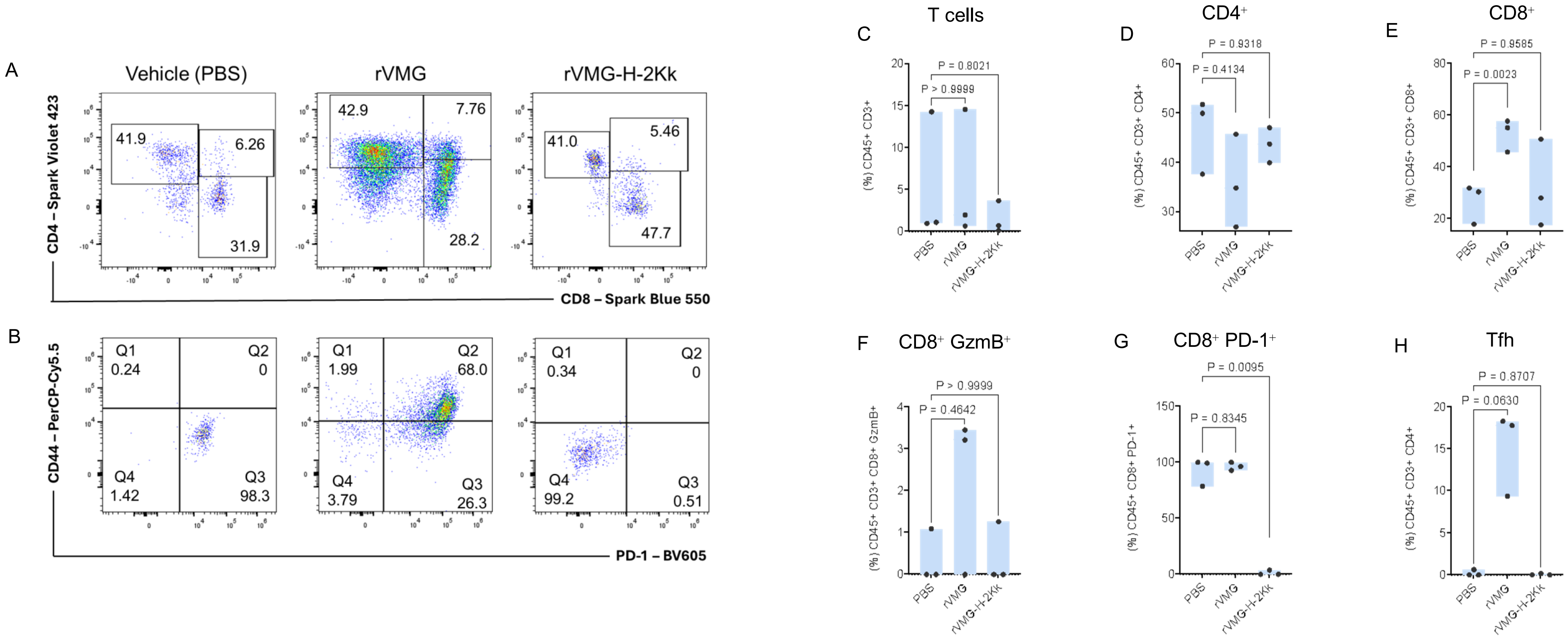

### Supplemental Fig. 3

A

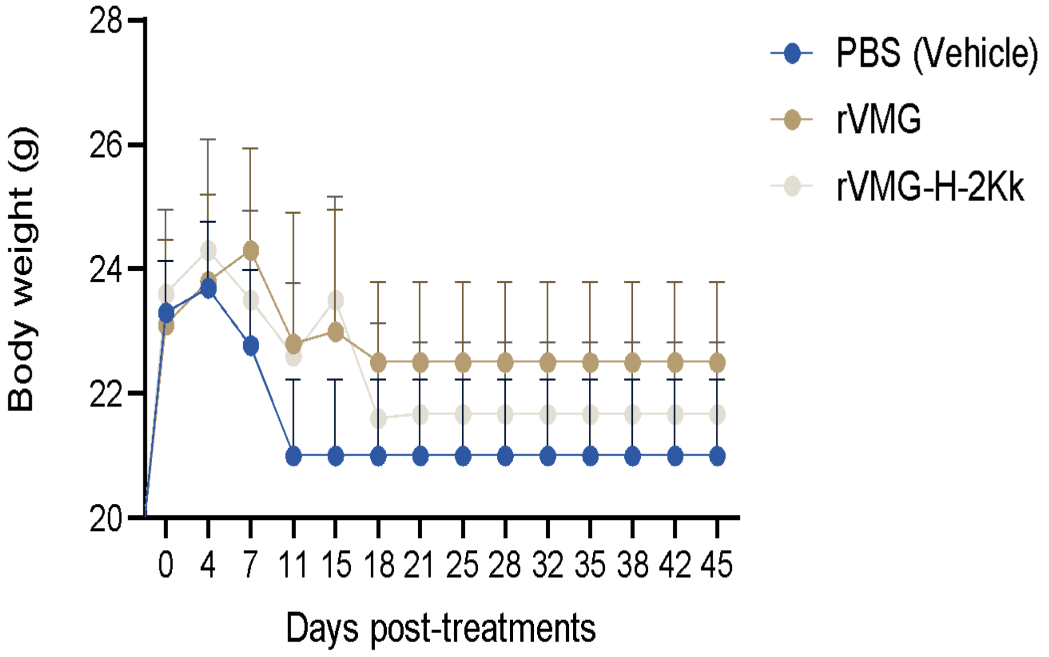


B

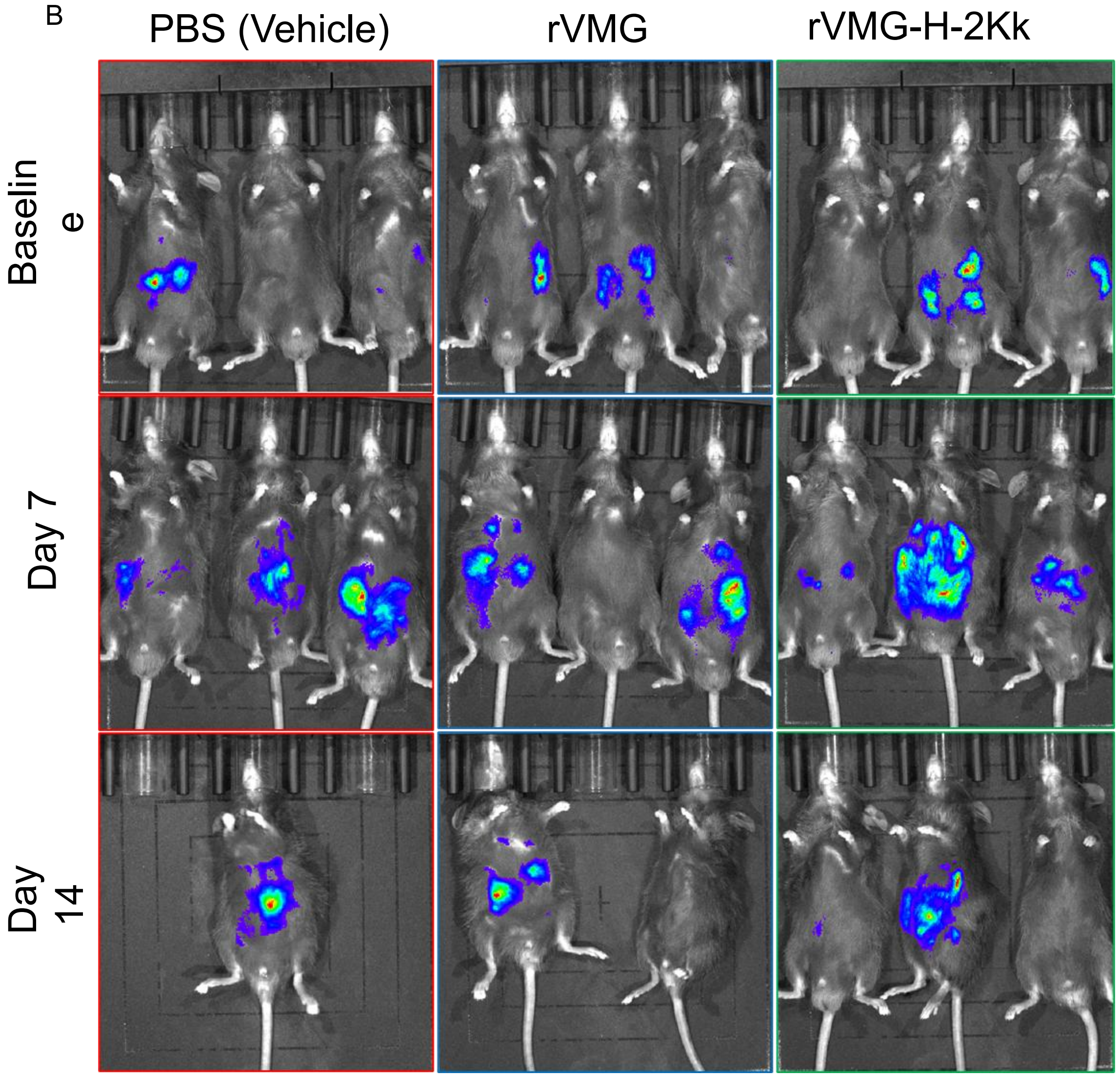

Supplemental Fig. 4

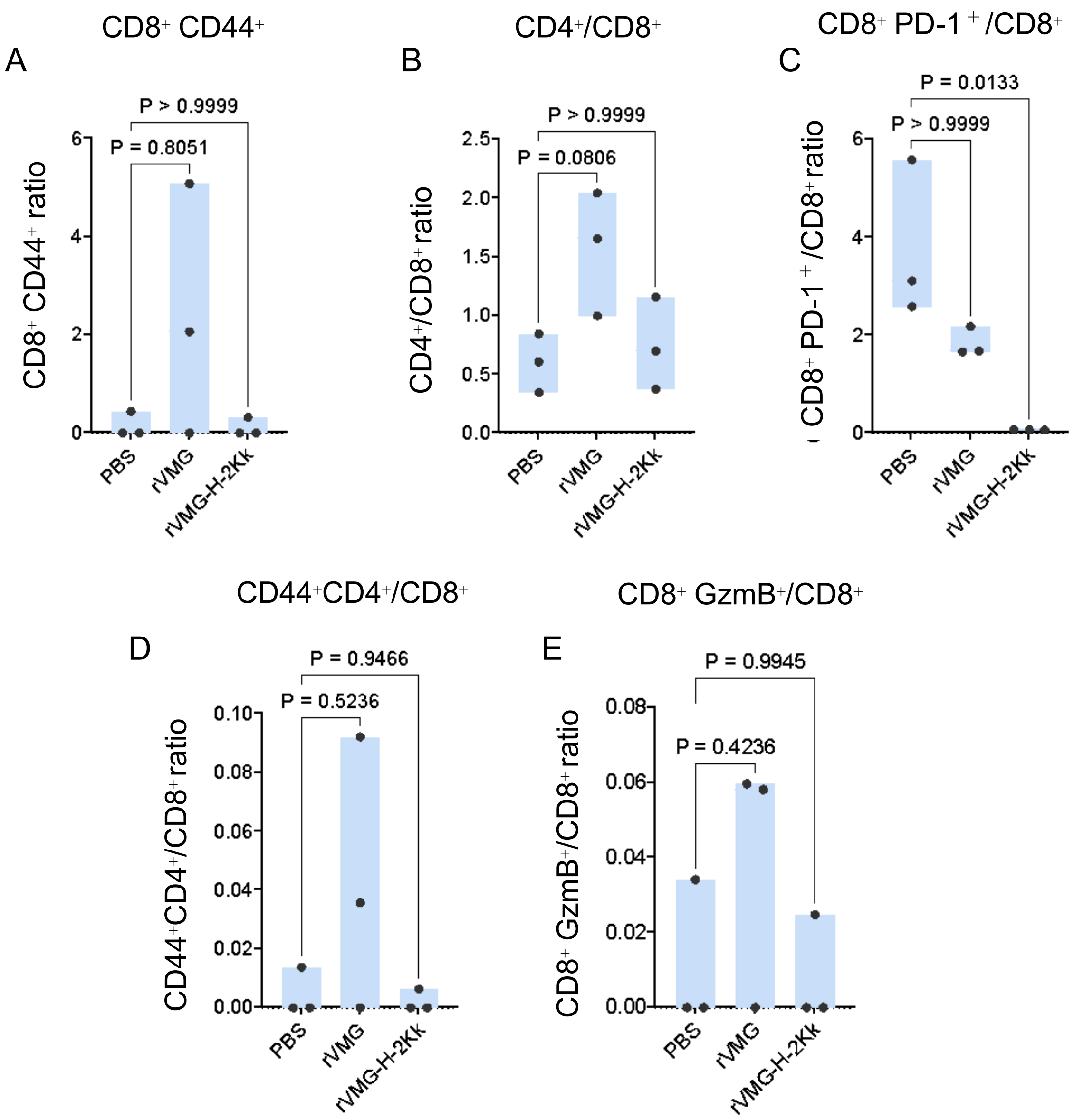

### Supplemental Fig. 5

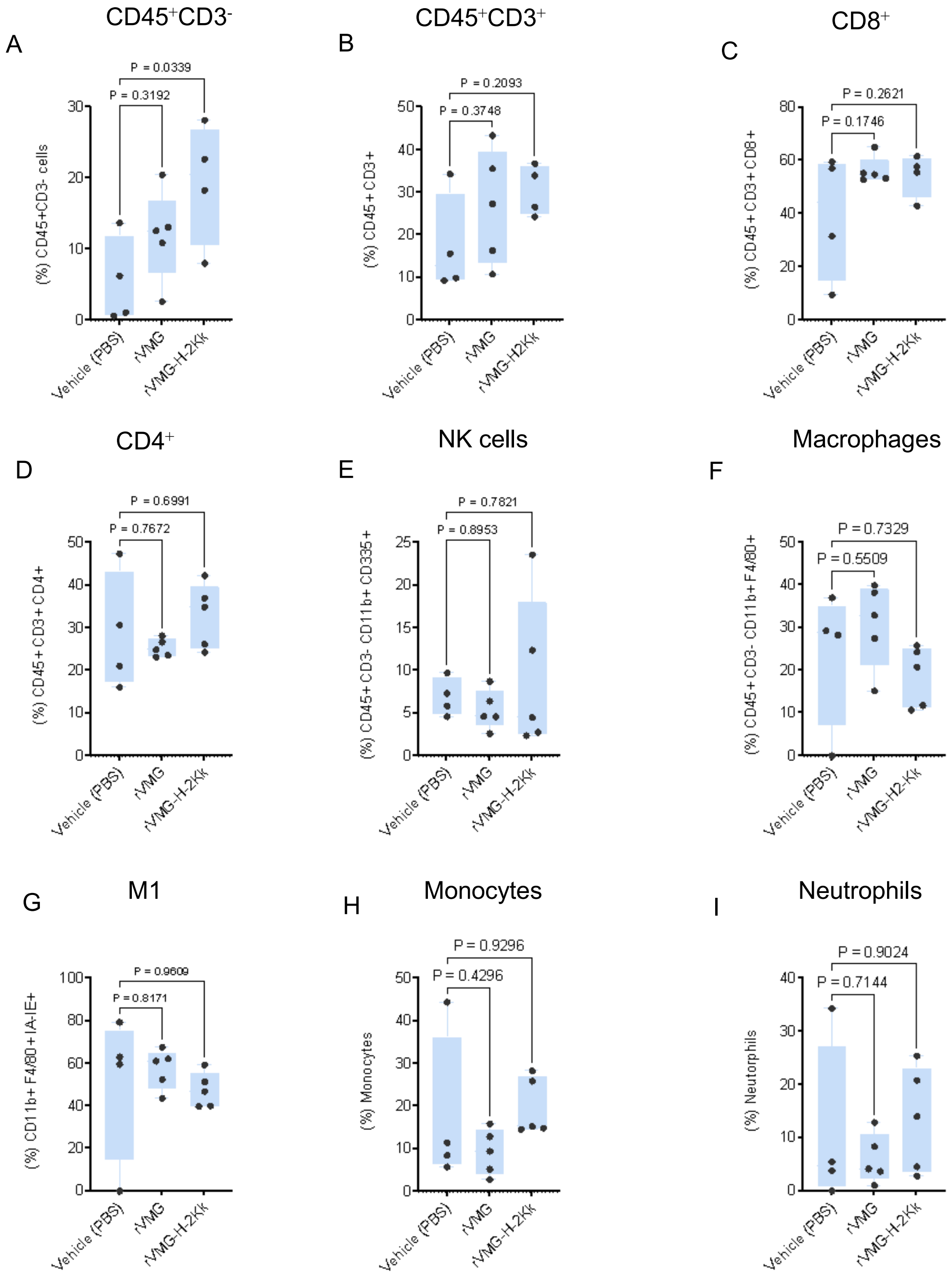

### Supplemental Fig. 6

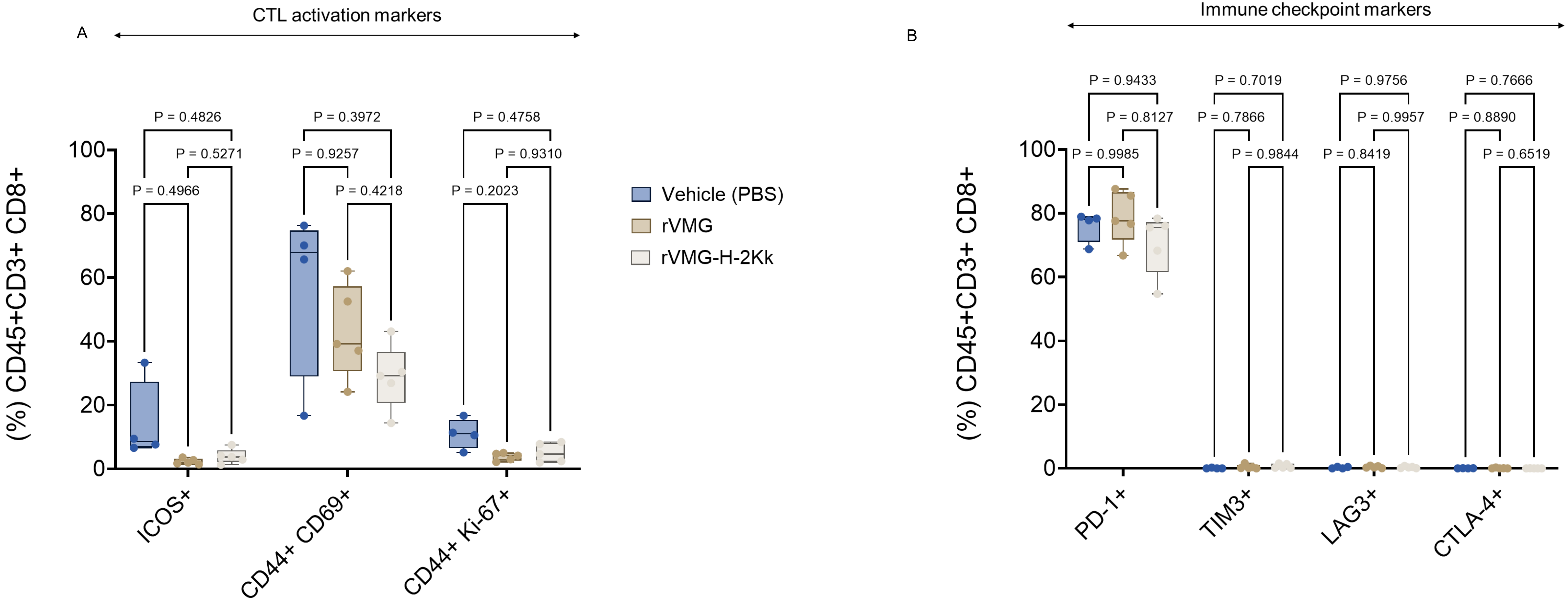

Supplemental Fig. 7

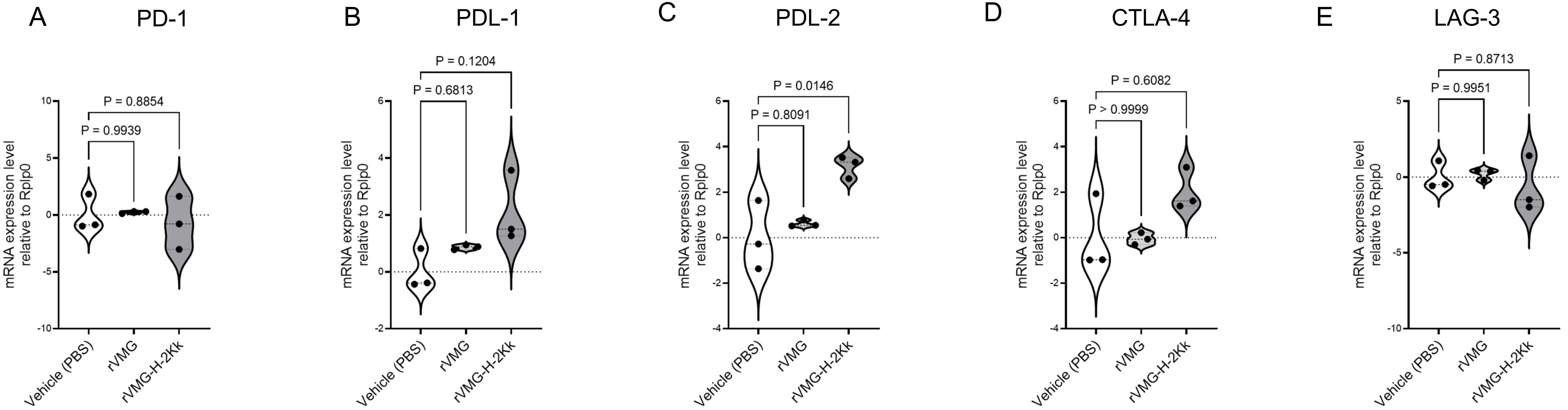

Supplemental Fig. 8

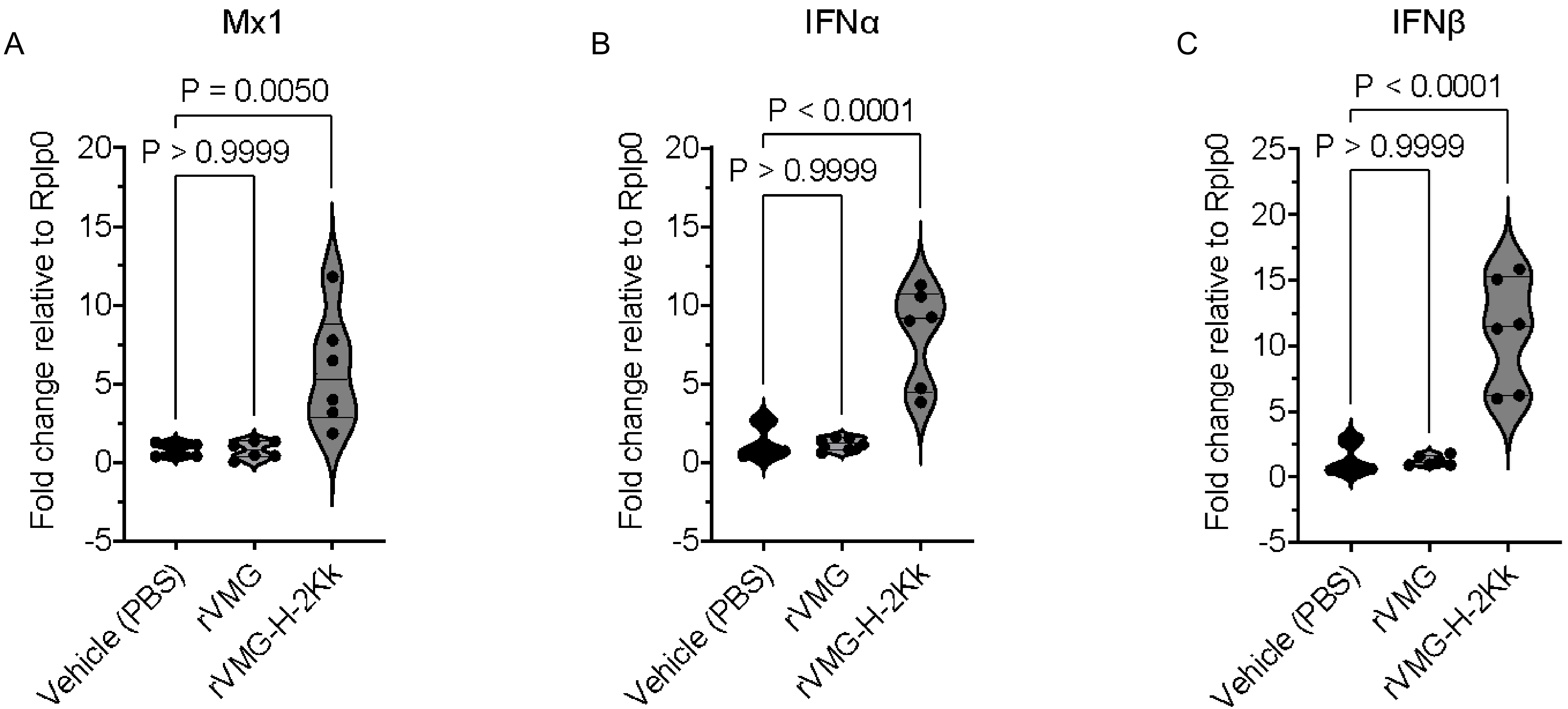

### Supplemental Fig. 9

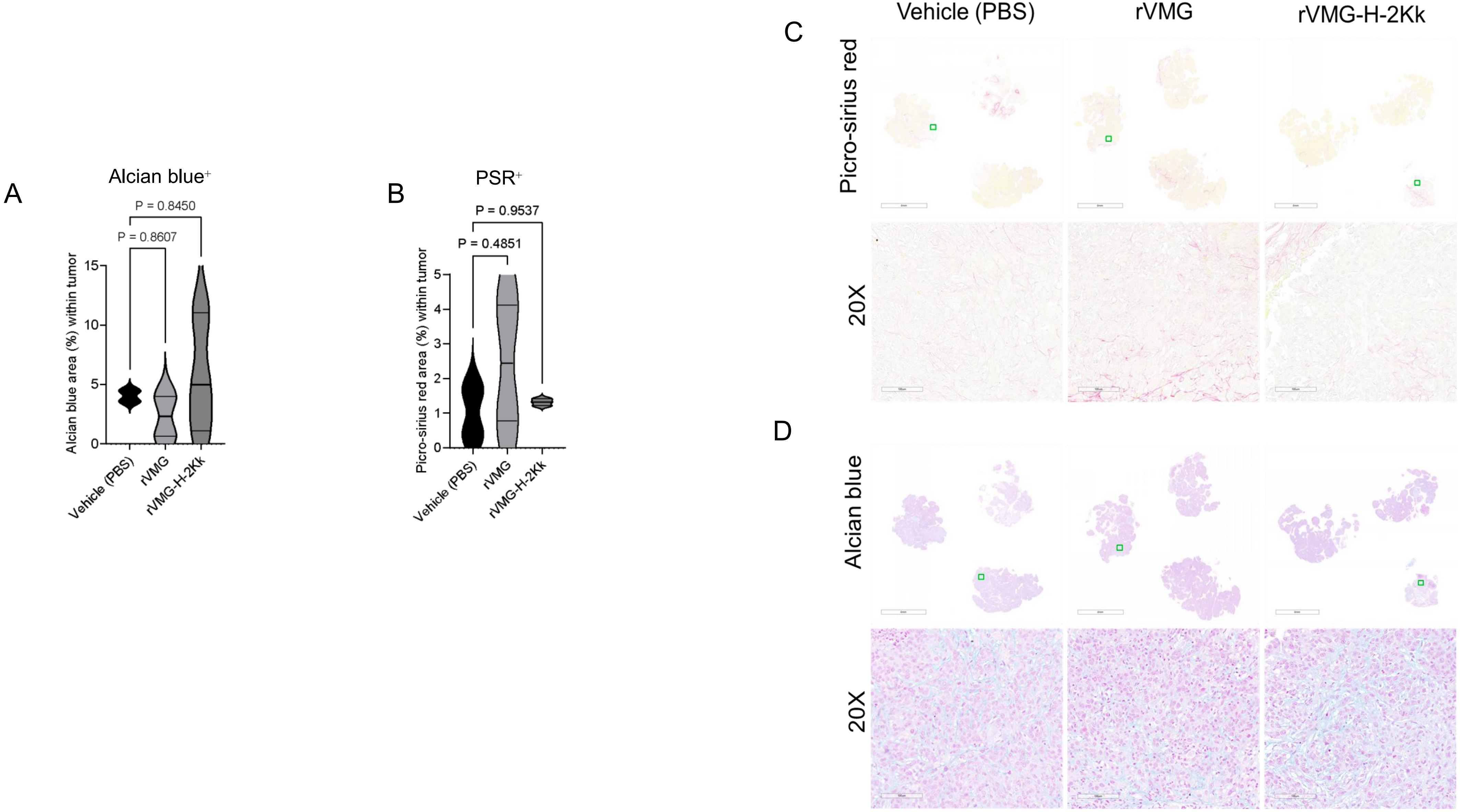

### Supplemental Figure 10

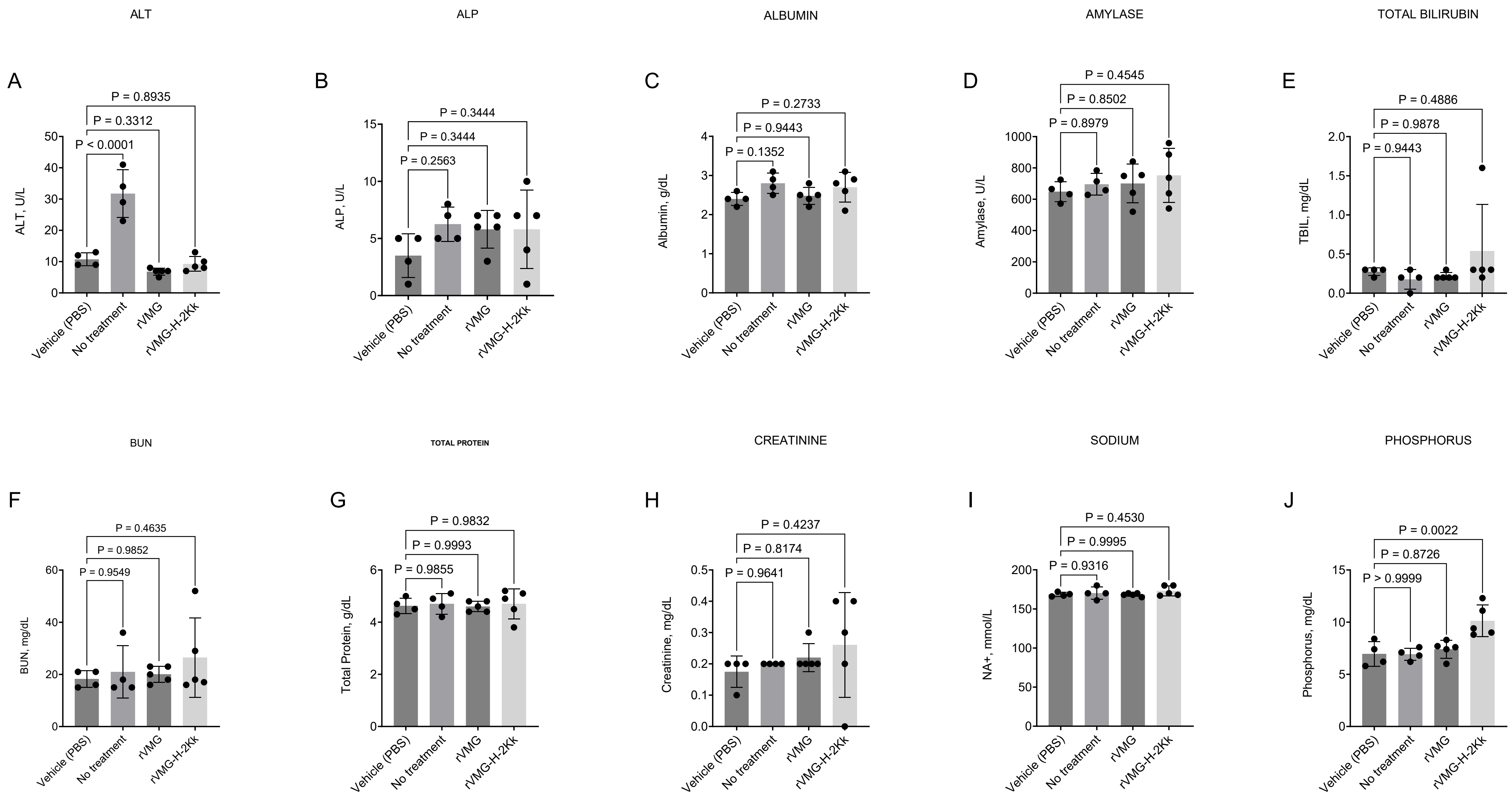

### Supplemental Fig.11

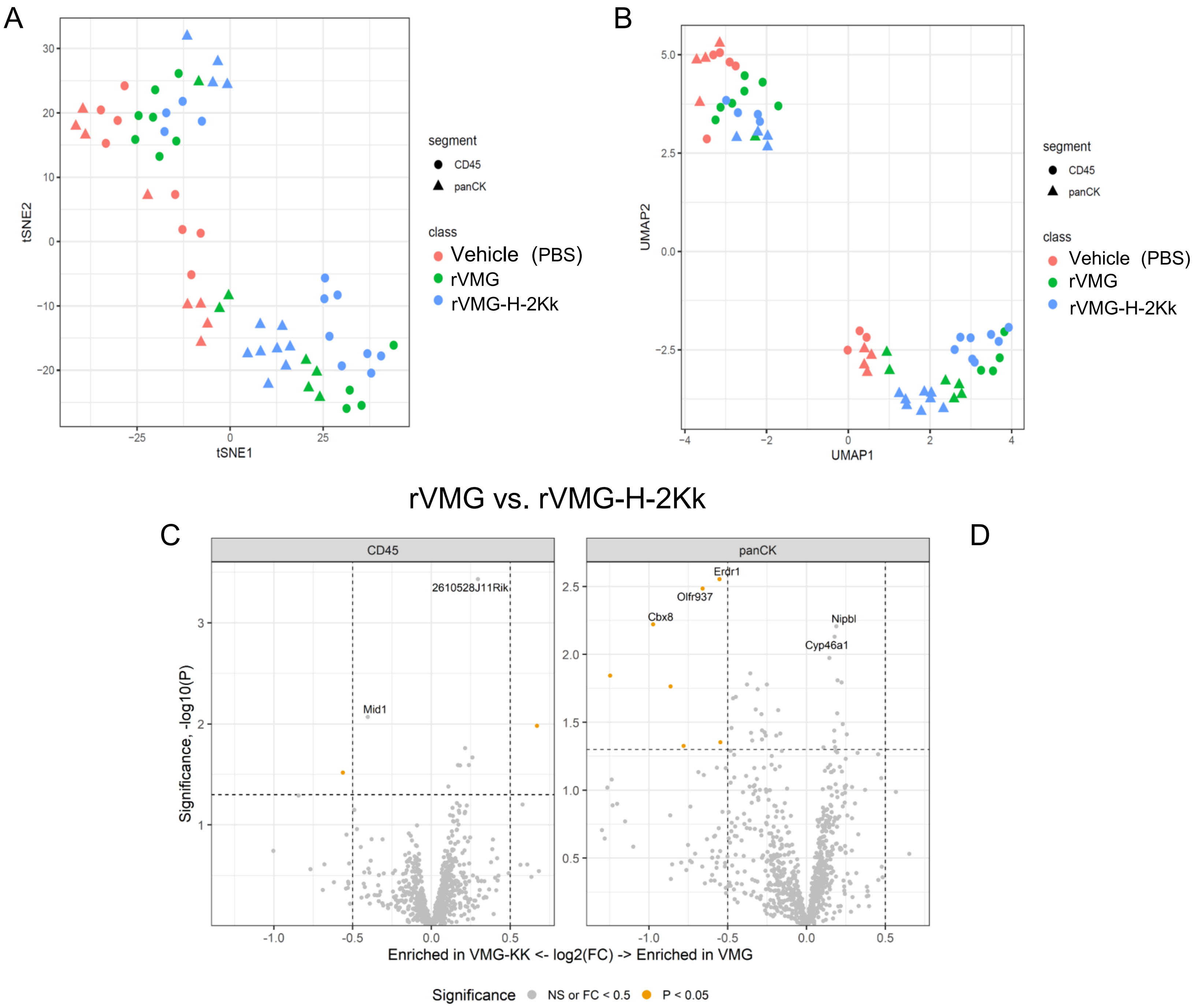

### Supplemental Fig. 12

Vehicle (PBS) vs. rVMG

A

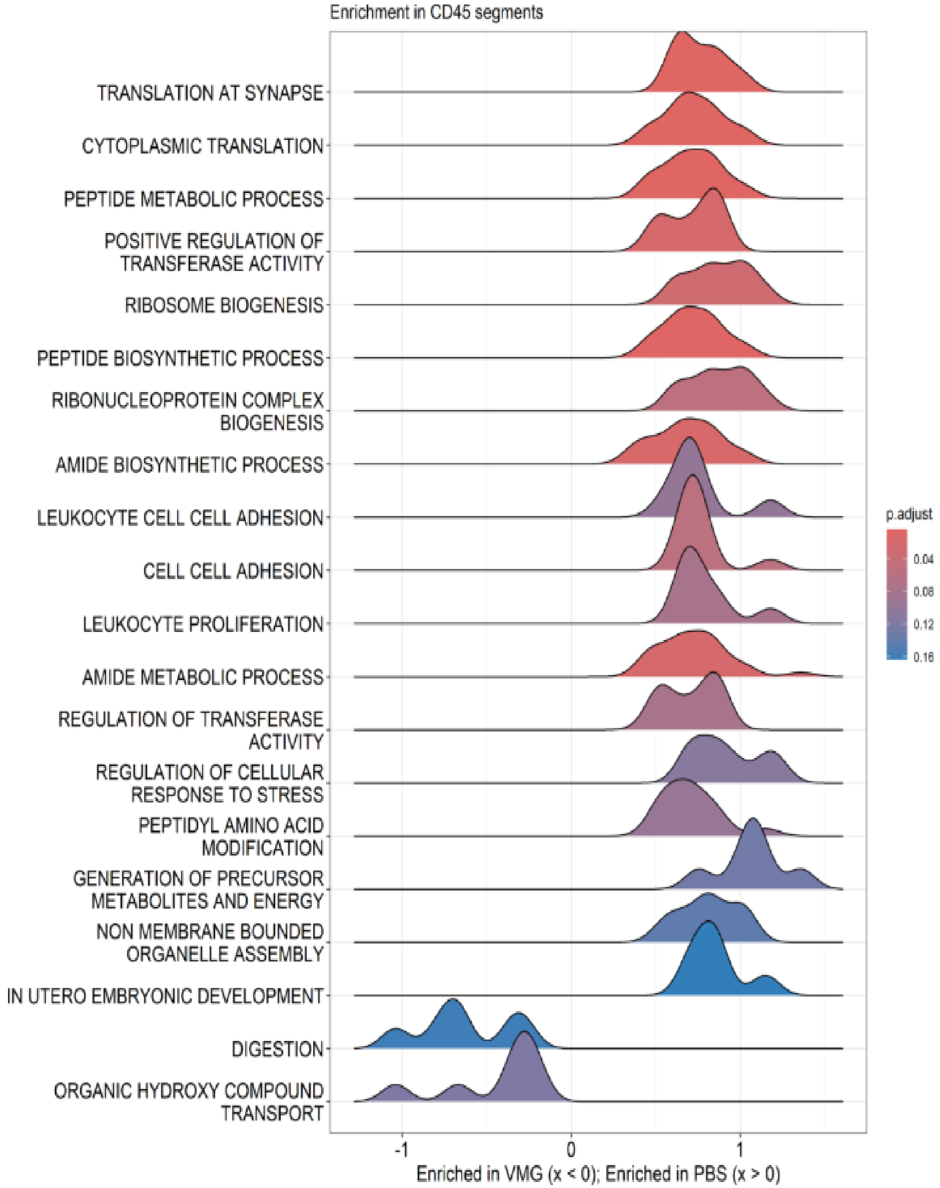


B

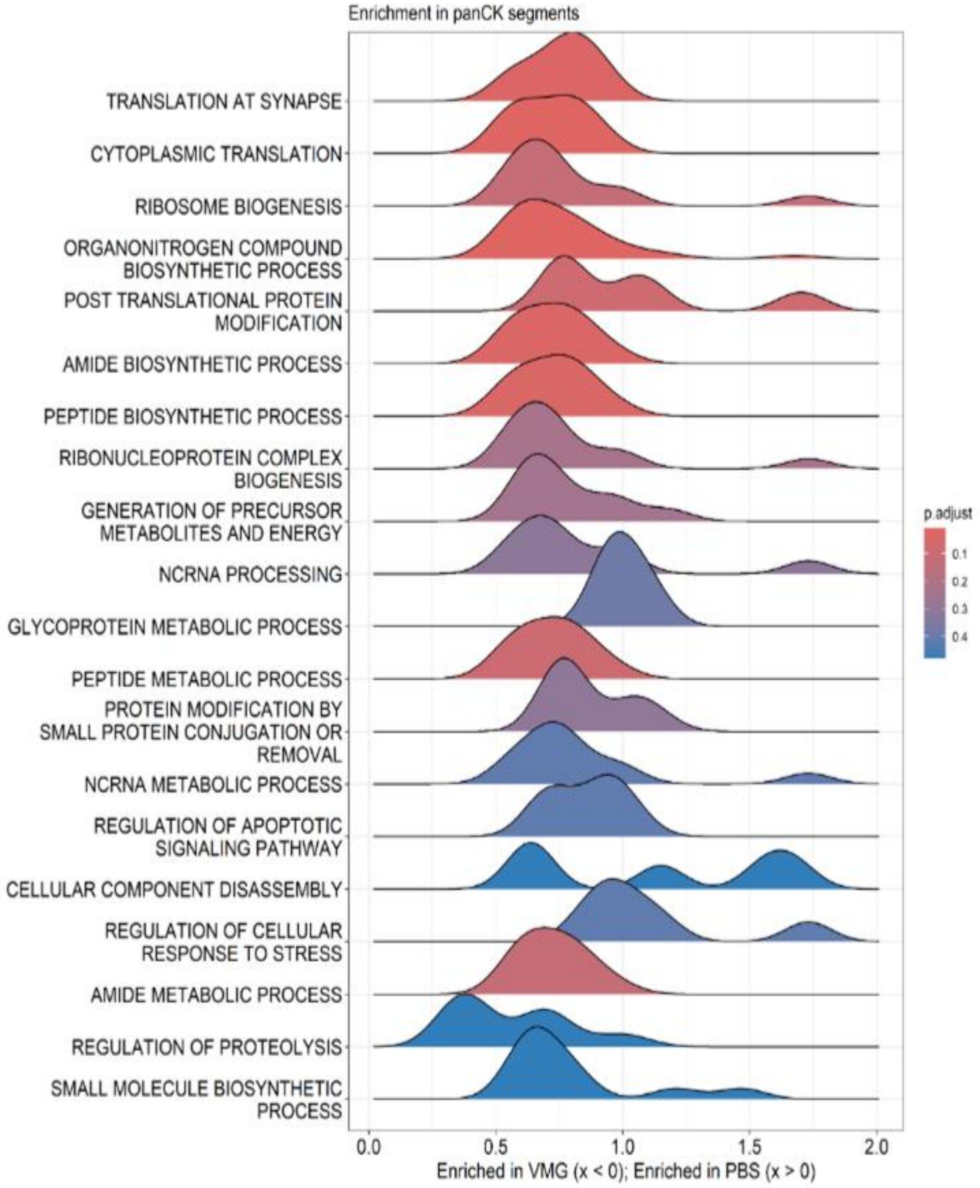

### Supplemental Fig. 13

Vehicle (PBS) vs. rVMG-H-2Kk

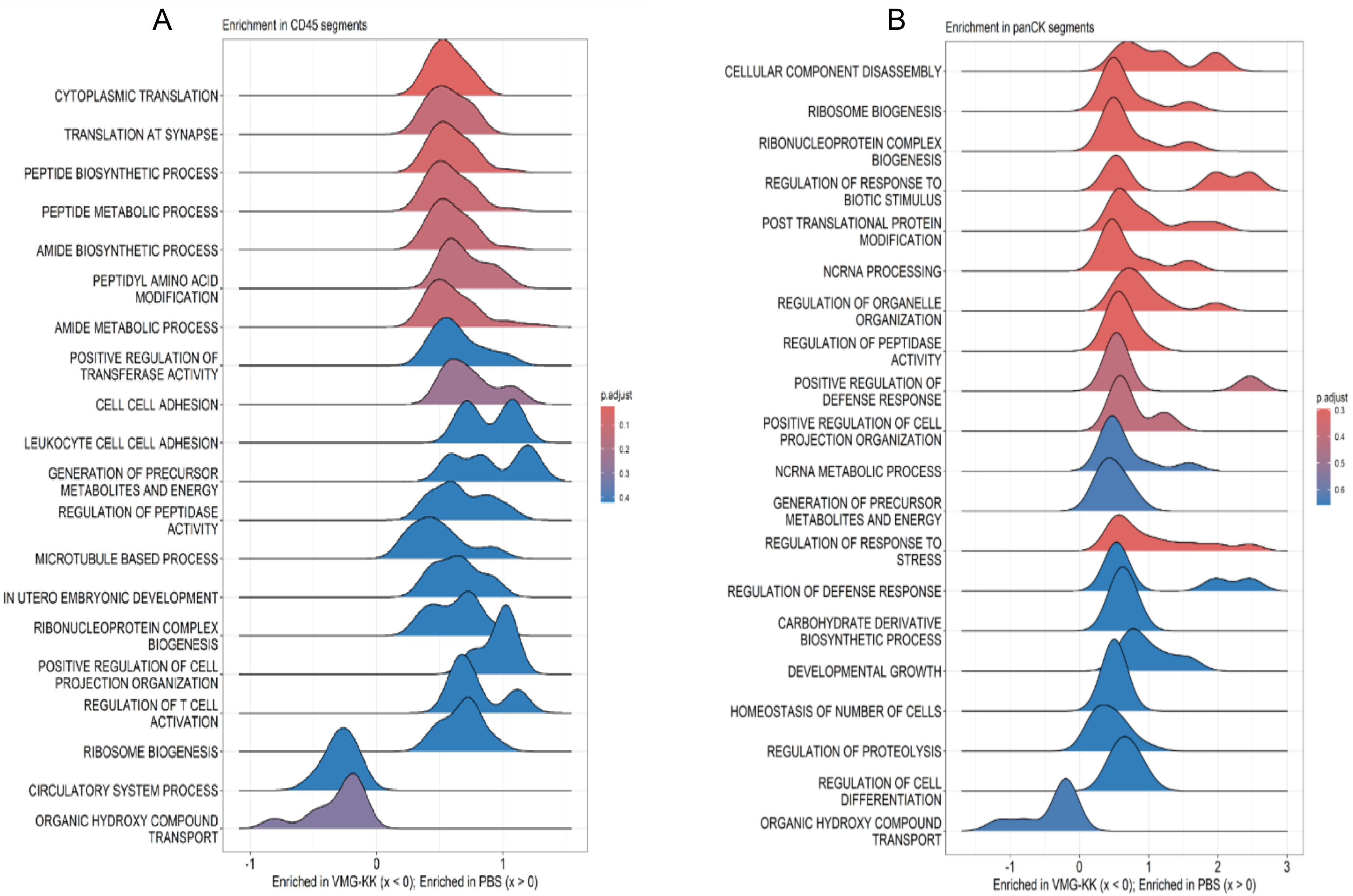

Supplemental Fig. 14

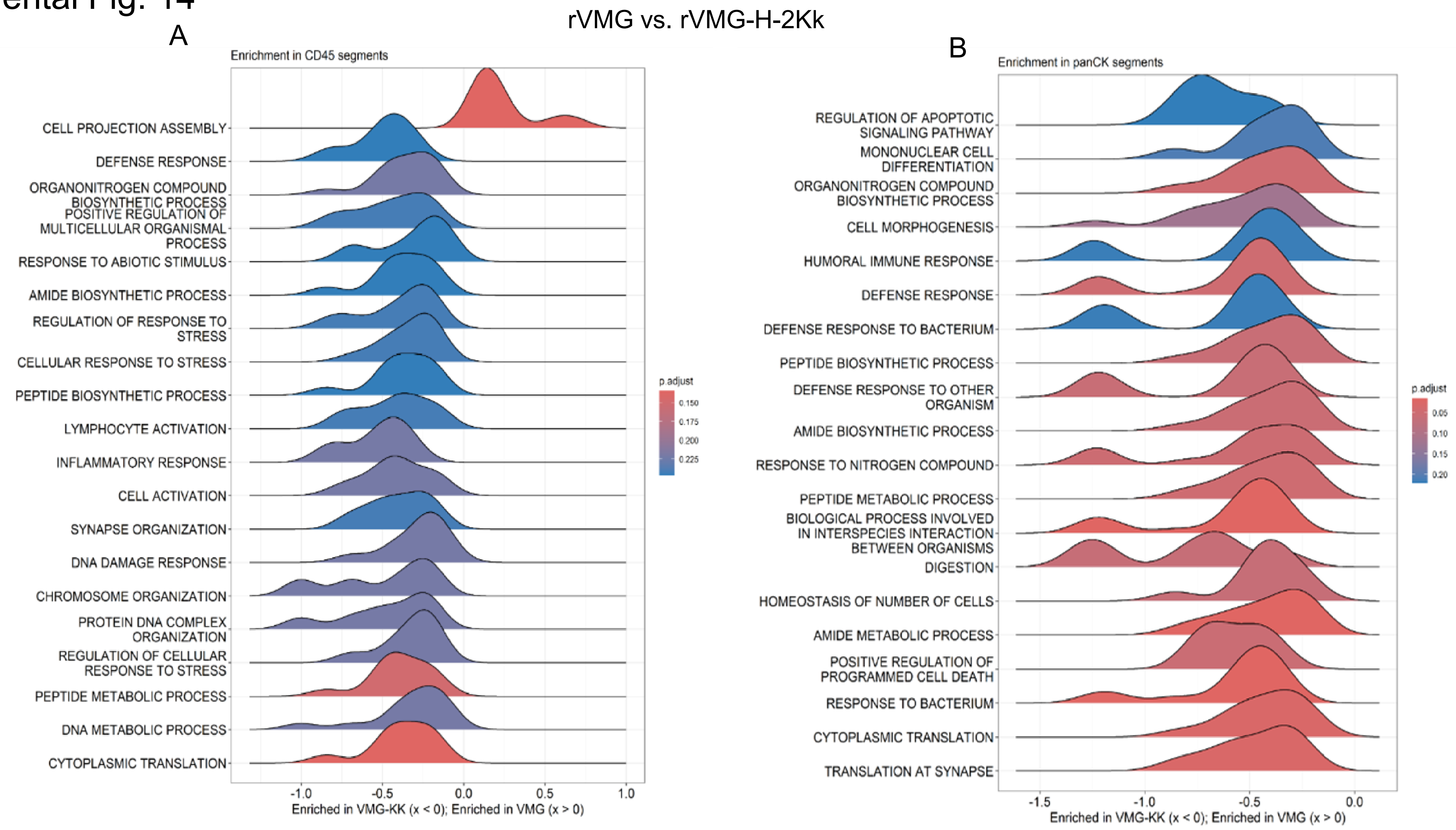

Supplemental Fig. 15

A

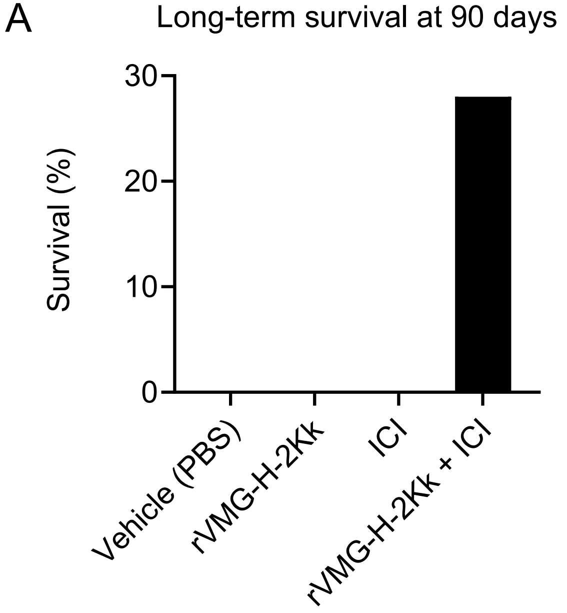


B

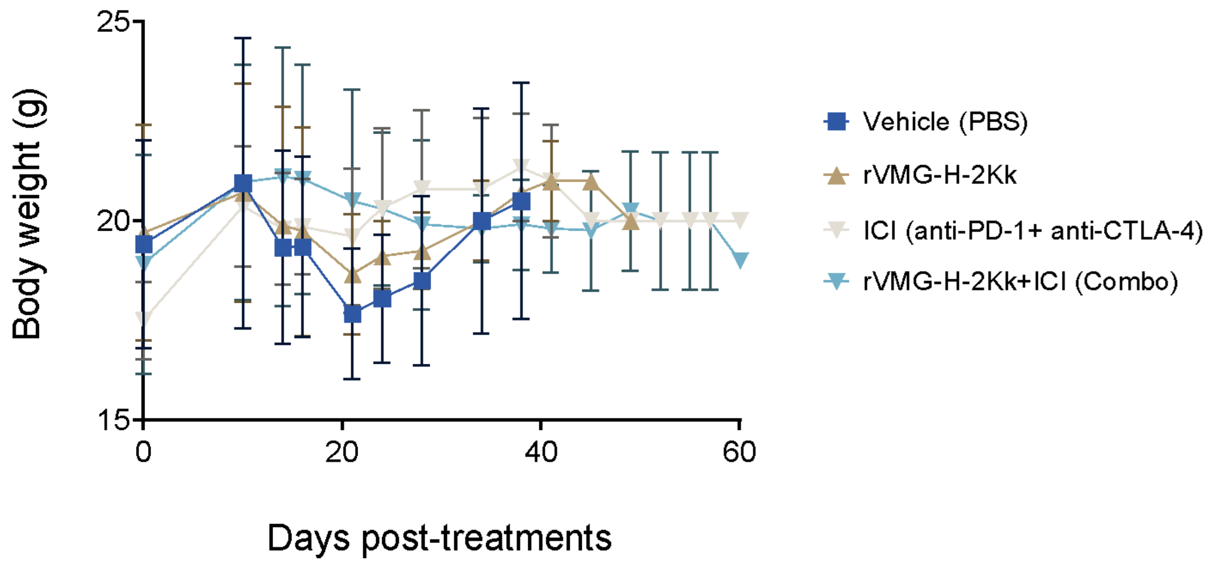

### Supplemental Fig. 16

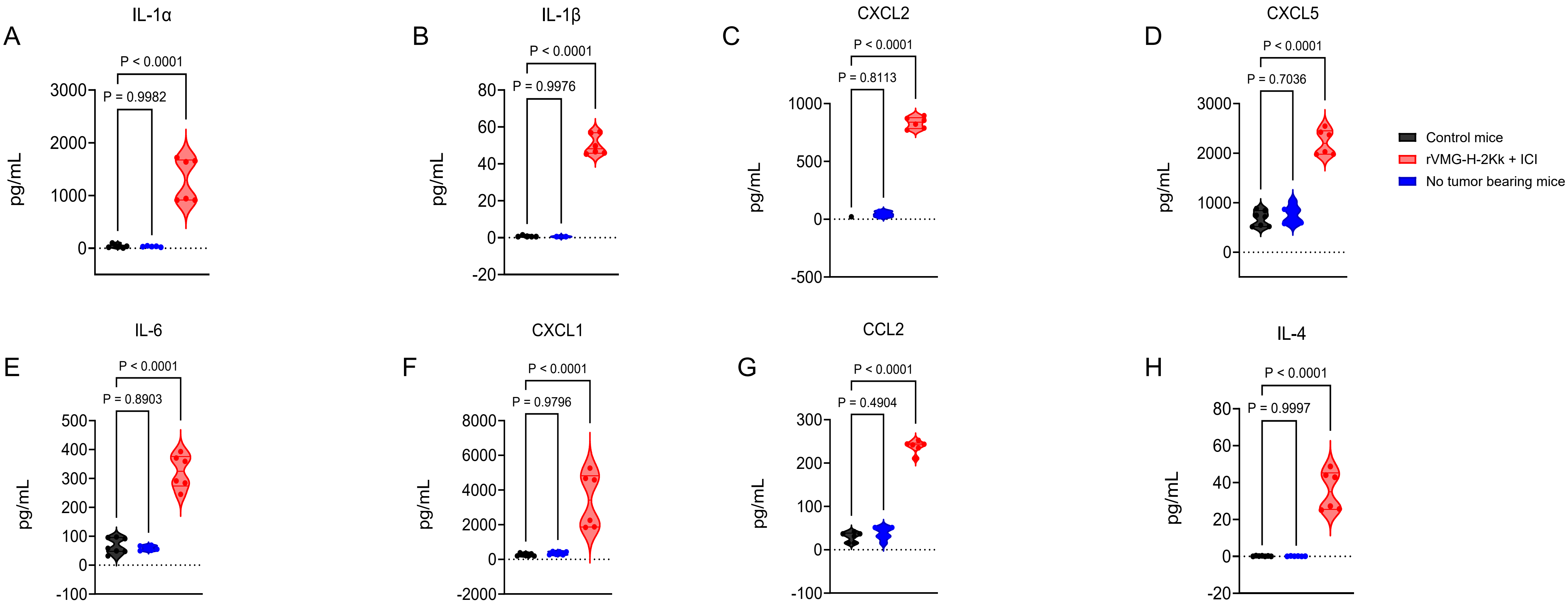

Supplemental Fig. 17

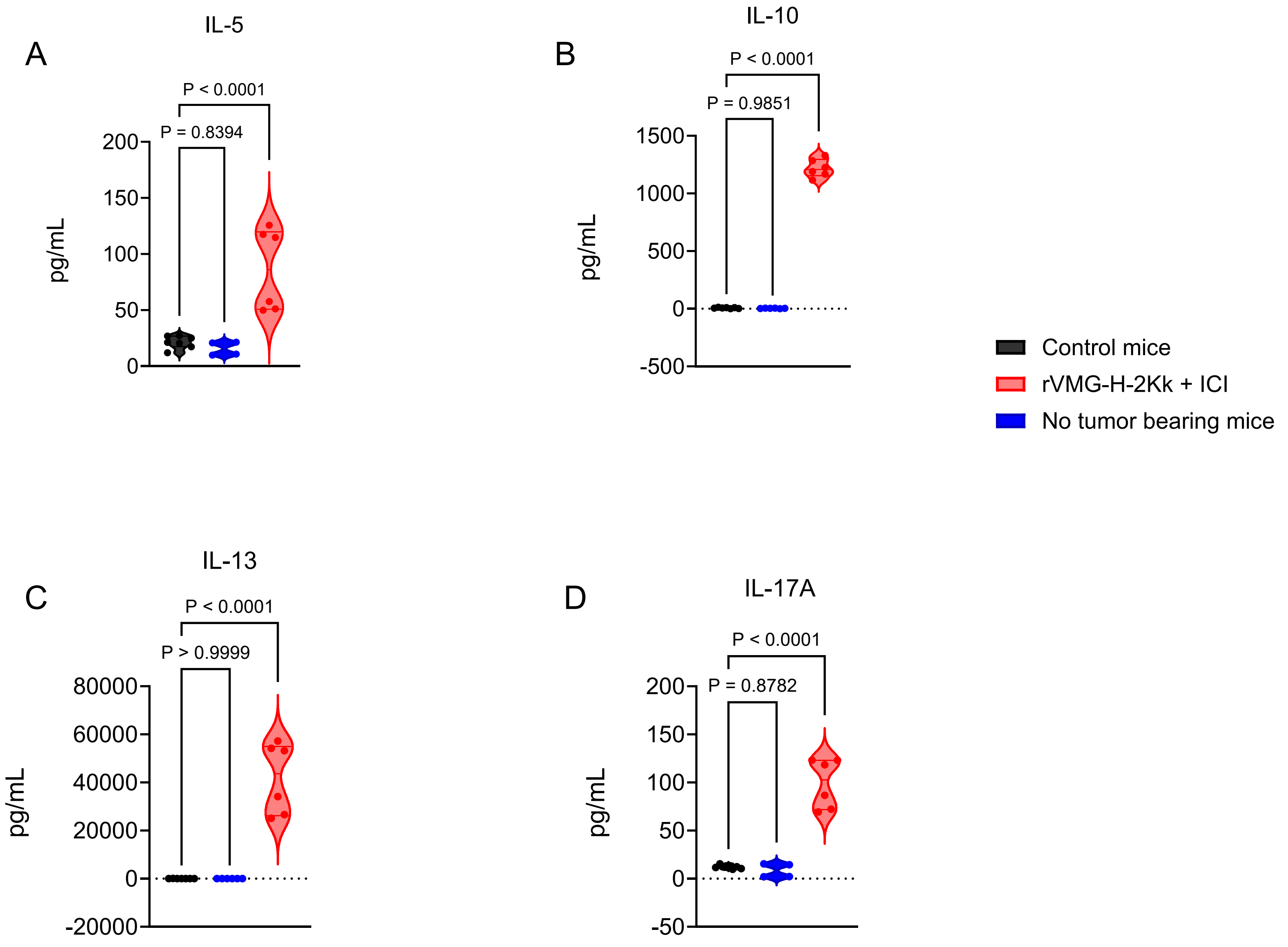

### Supplemental Table 1

**Table 1**: List of antibodies (IHC)

| Antibody | Company | Cat. No. | Type | Reactivity |
|---|---|---|---|---|
| CD8α (D4W2Z) | Cell Signaling | 98941 | Rabbit IgG mAb | Mouse |
| F4/80 | Cell Signaling | 70076 | Rabbit IgG mAb | Mouse |
| Granzyme B | Cell Signaling | 46890 | Rabbit IgG mAb | Human, Mouse |
| MHC II | Invitrogen | 14-5321-82 | Rat IgG2b mAb | Human, Mouse |
| FoxP3 | Cell Signaling | 12653 | Rabbit IgG mAb | Mouse |

#### Supplemental Table 2

| Antibody (Biolegend) | Clone | Cat. no | Fluorochrome |
|---|---|---|---|
| Live/Dead cells | - | 65-0865-14 | Zombie yellow |
| CD45 | 30-F11 | 103174 | APC-Fire 810 |
| CD3 | 17A2 | 100246 | PE-Dazzle 594 |
| CD4 (L3T4) | RM4-5 | 100578 | Spark Violet 423 |
| CD8a | 53-6.7 | 100780 | Spark Blue 550 |
| CD11b | M1/70 | 101243 | BV785 |
| CD11c | N418 | 117353 | BV510 |
| F4/80 | BM8 | 123137 | BV421 |
| CD335 (NKp46) | 29A1.4 | 137608 | APC |
| CD206 (MMR) | C068C2 | 141720 | PE-Cy7 |
| Ly6G | 1A8 | 127641 | BV650 |
| Ly6C | HK1.4 | 128024 | AF700 |
| I-A/I-E | M5/115.5.2 | 107618 | AF488 |
| CD279 (PD-1) | 29F.1A12 | 135220 | BV605 |
| CD44 | IM7 | 103032 | PcP-Cy5.5 |
| CD19 | 6D5 | 115558 | APC-Fire 750 |
| CD278 (ICOS) | C398.4A | 313558 | BV750 |
| CD366 (Tim-3) | QA20A20 | 165404 | PE |
| CD223 (LAG-3) | C9B7W | 125248 | PE/Fire 640 |
| CD152 (CTLA-4) | UC10-4B9 | 106335 | PE/Fire 810 |
| CD69 | H1.2F3 | 104558 | Spark NIR 685 |
| Ki-67 | 11F6 | 151227 | BV711 |

Supplemental Table 3. Mouse qPCR primer sequences

| Gene | Forward Primer (5'–3') | Reverse Primer (5'–3') |
|---|---|---|
| Rplp0 | CGCTTGTACCCATTGATGATG | TTATAACCCTGAAGTGCTCGAC |
| H-2Kk | GACACCCAGTTCGTCAGATT | TGTTTCTCCGCCTCATCTTC |
| H-2Kb | TGGCTCCTACACAGACAAGA | CTGTGTCTCCCTCTCCCAATA |
| B2M | ACAGTTCCACCCGCCTCACATT | TAGAAAGACCAGTCCTTGCTGAAG |
| Tap1 | GACTCCTTGCTCTCCACTCAGT | AACGCTGTCACCGTTCCAGGAT |
| Tapbp | TGGTCAGCGTATCCAGCACTCT | TTATGGGTGAGGACGGTCAGCA |
| VSV-N | TGTCTACCAAGGCCTCAAATC | CCTGCTTTCCCGATGTTTATTC |
| MX1 | TGGACATTGCTACCACAGAGGC | TTGCCTTCAGCACCTCTGTCCA |
| IFNα | TGCCCAGCAGATCAAGAAGG | TCAGGGGAAATTCCTGCACC |
| IFNβ | GCCTTTGCCATCCAAGAGATGC | ACACTGTCTGCTGGTGGAGTTC |
| PD-1 | CGGTTTCAAGGCATGGTCATTGG | TCAGAGTGTCGTCCTTGCTTCC |
| PD-L1 | CGCCTGCAGATAGTTCCCAA | TAAGGTCCTCCTCTCCTGCC |
| PD-L2 | CTGGGACTACAAGTACCTGACG | CTCTAGCCTGGCAGGTAAGCTG |
| CTLA-4 | GTACCTCTGCAAGGTGGAACTC | CCAAAGGAGGAAGTCAGAATCCG |
| LAG-3 | CTCCATCACGTACAACCTCAAGG | GGAGTCCACTTGGCAATGAGCA |

Supplemental Table 4

| Group | Median Survival (days) | Long-term Survivors at 90 Days |
|---|---|---|
| Vehicle (PBS) | ~20 | 0% |
| rVMG-H-2Kk | ~28–30 | 0% |
| ICI (anti-PD-1 + anti-CTLA-4) | ~41–45 | 0% |
| rVMG-H-2Kk + ICI (Combo) | >90 (Not Reached) | 27% |
